# A cold-inducible phospholipid–protein interaction in brown fat mitochondria optimizes thermogenic capacity

**DOI:** 10.1101/2025.05.15.654206

**Authors:** Yuta Shimanaka, Marcus J. Tol, Christian Rocha-Roa, Matthew J. Jellinek, Liujuan Cui, Aaron Bender, Alexander H. Bedard, Madeleine G. Milner, Bruno Melillo, Bassem M Shoucri, Adrian Wong, Kevin J. Williams, Linsey Stiles, Michael Shum, Thomas A. Weston, Whitaker Cohn, Julian P. Whitelegge, Itay Budin, Ambre M. Bertholet, Benjamin F. Cravatt, Stephen G. Young, Sander M. Houten, Carmen Argmann, David A. Ford, Marc Liesa, Orian S. Shirihai, Stefano Vanni, Peter Tontonoz

**Author notes:** These authors contributed equally to this work (Yuta Shimanaka, Marcus J. Tol).

## Abstract

Cold stress elicits dynamic remodeling of the mitochondrial lipidome in brown adipose tissue (BAT), marked by an increase in arachidonoyl-phosphatidylethanolamine (AA-PE). However, the function of membrane lipid rewiring in thermoregulatory physiology has been a longstanding mystery. Here, we identify LPCAT3 as a cold-regulated O-acyltransferase driving the highly selective accrual of AA-PE in BAT mitochondria. Lipid-based proteomics, molecular dynamics simulations, and bioenergetic analyses reveal that AA-PE partitions at the COX4I1 interface of the Cytochrome *c* oxidase complex, enhancing electron transport chain (ETC) efficiency. Accordingly, fat-specific *Lpcat3*-knockout mice have defects in respiratory-dependent BAT thermogenesis and cold tolerance, despite intact β-adrenergic signaling and UCP1 function. Under cold acclimation, *Lpcat3^−/−^*BAT exhibits ETC dysfunction and activation of the integrated stress-response. Thus, our study illuminates a cold-regulated lipid–protein interaction as a gating factor in UCP1-dependent thermogenesis.

---

Adipose tissue is a metabolic and endocrine organ that plays a critical role in systemic energy balance. White adipose tissue (WAT) acts as a key energy reservoir, while mitochondrial-dense BAT specializes in fuel combustion and is responsible for adaptive thermogenesis (*1, 2*). This heat-generating capacity is conferred in part by uncoupling protein 1 (UCP1), a brown fat-specific LCFA^−^/H^+^ symporter that enables futile proton cycling across the inner mitochondrial membrane (IMM) (*3, 4*). In addition, certain WAT depots can undergo “browning” in response to cold temperatures, giving rise to distinct subpopulations of beige adipocytes that engage in UCP1-dependent and -independent futile cycles (*5–7*). In humans, BAT was long appreciated to be present in newborns but to disappear by adulthood (*8, 9*). The discovery of brown/beige adipocytes in adult humans has rekindled interest in identifying factors that control and/or support UCP1-dependent uncoupled respiration (*10–14*).

Substantial progress has been made in understanding the pathways that control BAT identity and UCP1 function (*15*). One key unresolved question is how thermal stress converges on the membrane biophysical properties of BAT, and whether such adaptations contribute to its heat-generating capacity. Thermoregulatory control over the brown fat mitochondrial phospholipidome was first documented in the 1970s (*16–18*). Cold exposure is accompanied by an overall increase in phosphatidylcholine (PC), PE, and cardiolipin (CL) abundance in brown adipocytes (*17, 19, 20*). Conically shaped PE and CL are enriched in the IMM where they generate the negative curvature for cristae folding and ETC assembly (*21–26*). Interestingly, the BAT phospholipidome also has been reported to undergo qualitative changes (*17, 27*). These include a cold-inducible enrichment in polyunsaturated fatty acyl (PUFA) groups, such as those derived from linoleic (18:2*n*-6) and arachidonic acid (20:4*n*-6). Recent advances in lipidomics technology have revealed that thermal stress selectively enriches arachidonoyl-PE species in brown fat mitochondria (*19, 28*). However, the functional role of cold-induced membrane lipidome remodeling in thermoregulation has remained elusive due to the lack of experimental tools for manipulating acyl-chain composition *in vivo*.

Early studies using essential fatty acid-deficient diets implied a requirement for PUFAs during non-shivering thermogenesis and hibernation (*29–31*). BAT-specific deletion of PE and CL biosynthetic enzymes has been shown to cause global deficits in adrenergic signaling, thermogenic gene programs, and/or UCP1 activity (*20, 28*). More recently, elegant work has suggested that cold-induced membrane lipid remodeling elicits a conformational switch of BAT respiratory supercomplexes (SCs), enhancing catalytic efficiency (*32*). However, none of these studies have specifically addressed a requirement for AA-PE nor any other specific phospholipid species in thermoregulatory physiology. Polyunsaturated phospholipids are generated in the ER via sequential deacylation–reacylation reactions, known as Lands’ cycle (*33*). This dynamic process yields membrane acyl-chain asymmetry that varies depending on cell/tissue type and physiological state (*34, 35*). In this study, we identify LPCAT3 as the long-sought O-acyltransferase required for the thermal stress-induced accumulation of AA-PE in BAT mitochondria. We further uncover a molecular basis for the effects of AA-PE on respiratory-dependent BAT thermogenesis.

## Cold-inducible AA-PE in thermogenic fat requires LPCAT3

We studied the consequence of thermal stress on the phospholipidome of brown/beige fat depots by acclimating wild-type mice to either 5 ℃ or thermoneutrality (TN, 30 ℃), which minimizes β-adrenergic tone. Shotgun-lipidomic profiling of BAT and isolated brown fat mitochondria revealed enrichment of AA-PE species, most notably 18:0_20:4-PE, in response to cold (Fig. 1, A and B). Cold-induced browning of inguinal WAT (iWAT) was also associated with a marked rise in AA-PE levels (Fig. 1C). This lipidomic rewiring was specific to thermogenic fat, as no such change was found in quadriceps muscle (Fig. 1D). The BAT membrane lipidome is shaped by a subset of cold-inducible genes crucial for fatty acid elongation and glycerophospholipid remodeling (*36*). We focused our attention on LPCAT3, an ER membrane-bound O-acyltransferase that selectively enriches *n*-6 PUFAs in phospholipids (*37–39*). *Lpcat3* was the most abundantly expressed PC/PE remodeling enzyme in BAT (Fig. 1, E and F), comparable to its expression in the liver, intestine, and kidney. In addition, adipocyte *Lpcat3* mRNA levels were highly upregulated in response to thermal stress or β3-adrenoreceptor agonism (Fig. 1, G and H).

**Fig. 1.**
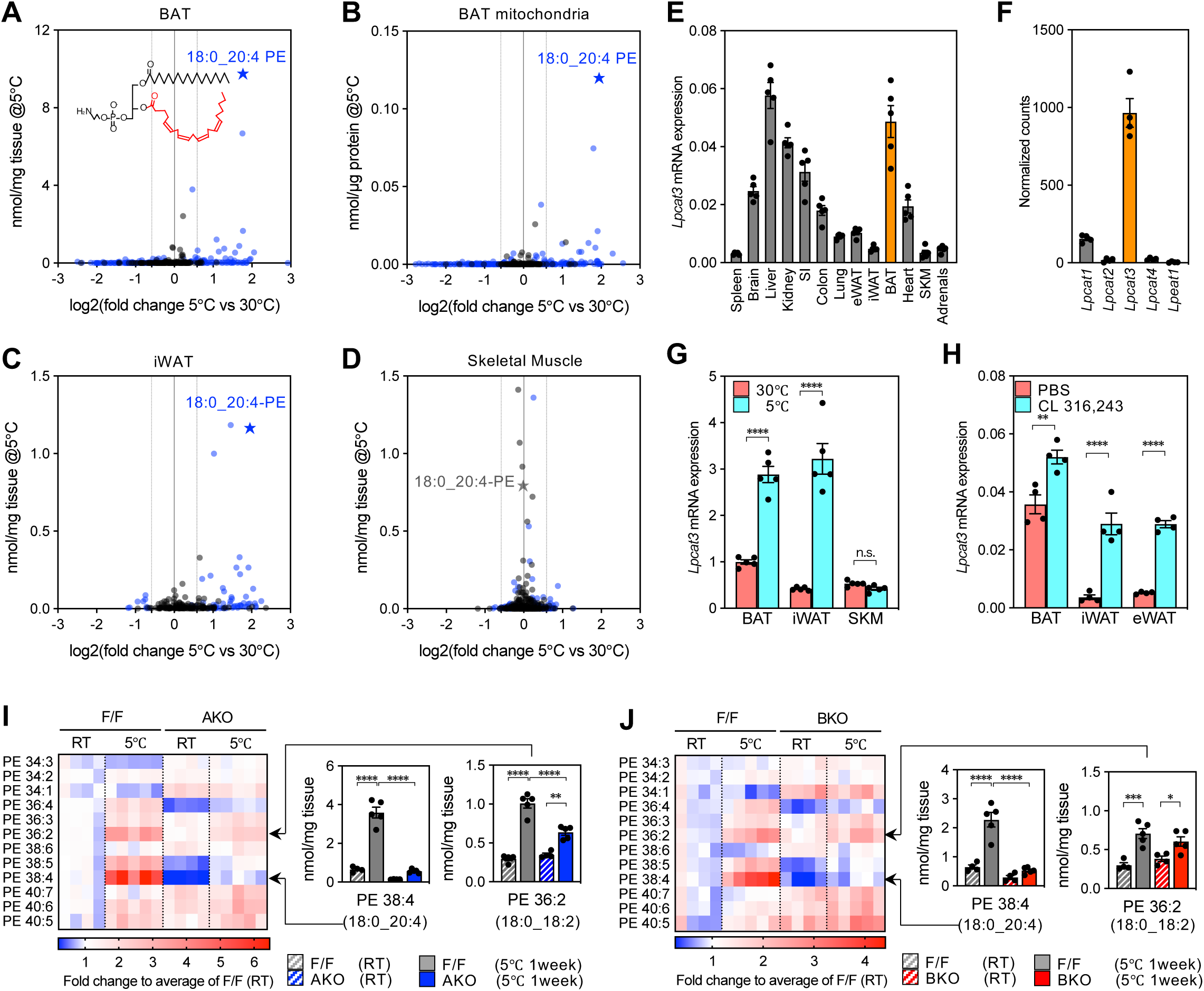
Identification of the O-acyltransferase for cold-inducible PE remodeling in thermogenic fat. (**A-D**) Shotgun-lipidomic analysis of (**A**) BAT, (**B**) purified BAT mitochondria, (**C**) iWAT, and (**D**) quadriceps muscle from 10-week-old wild-type mice housed at 30℃ or 5℃ for 7 days (*n* = 4, 4). Significantly changed (*p* < 0.05) lipid species after cold exposure are highlighted (blue) in volcano plots. (**E**) qPCR analysis of *Lpcat3* transcript levels in a panel of metabolic tissues from wild-type mice housed at 23℃ (*n* = 5). (**F**) *Lpcat1-4* transcript levels (normalized counts) in BAT from wild-type mice (*n* = 4). Data values were from a recently published RNA-seq dataset (GSE294663). (**G**) qPCR analysis of *Lpcat3* transcript levels in BAT, iWAT and quadriceps muscle (SKM) from 10-week-old wild-type mice housed at either 30℃, 23℃, or 5℃ for 7 days (*n* = 5/group). (**H**) qPCR analysis of *Lpcat3* transcript levels in BAT, iWAT, and eWAT from 10-week-old wild-type mice injected daily with PBS or CL-316,243 for 4 days (*n* = 4, 4). (**I, J**) LC-MS/MS analysis of abundant PE molecular species in BAT from (**I**) *Lpcat3*^AKO^ mice and F/F controls, or (**J**) *Lpcat3*^BKO^ mice and F/F controls housed at 23℃ or 5℃ for 7 days (*n* = 4–5/group). 38:4-PE and 36:2-PE subspecies are depicted in individual bar graphs. Data are presented as mean ± SEM. \**p*<0.05, \*\**p*<0.01, \*\*\**p*<0.001, \*\*\*\**p*<0.0001 by two-sided Welch’s *t*-test (G, H), or one-way ANOVA with Tukey’s multiple comparisons test (I, J).

To assess whether LPCAT3 activity is required for cold-induced remodeling of the BAT membrane lipidome, we employed pan-adipose *Lpcat3*-KO (*Adipoq*-Cre; *Lpcat3*^F/F^, hereafter *Lpcat3*^AKO^) and BAT-specific *Lpcat3*-KO (*Ucp1*-Cre; *Lpcat3*^F/F^, hereafter *Lpcat3*^BKO^) models (*40*) (fig. S1, A and B). *Lpcat3*^AKO^ and *Lpcat3*^BKO^ mice acclimated to cold in a manner comparable to their floxed (F/F) littermate controls and maintained euthermia when transferred from room temperature (23℃) to 5℃ (fig. S1, C to E). Notably, thermal stress-dependent AA-PE accrual was almost entirely abolished in *Lpcat3^−/−^* BAT (Fig. 1, I and J). This specific deficit was associated with a compensatory rise in monounsaturated fatty acid-(MUFA) and docosahexaenoyl-chains (*e.g.*, 34:1, 40:6, 40:7-PE). On the other hand, loss of LPCAT3 had a modest effect on AA-PC (fig. S1, F and G). AA-PE depletion was recapitulated in isolated brown fat mitochondria from *Lpcat3*^BKO^ mice, with no overt changes in the acyl chain profiles of other phospholipid classes (fig. S1, H and I). These data suggest that LPCAT3 is both necessary and sufficient for the cold-induced accrual of AA-PE in thermogenic fat.

## LPCAT3 activity enhances UCP1-dependent thermogenesis in brown adipocytes

We next explored the cell-intrinsic role of AA-PE in ETC functionality and respiratory-dependent thermogenesis using primary brown adipocytes (pBAs). We engineered an inducible *Lpcat3*-KO (IKO) system by crossing *Lpcat3*^F/F^ mice with the R26-*Cre*^ERT2^ model. The BAT-derived *Lpcat3*^F/F^, *Cre*^ERT2^ preadipocyte cell fraction was treated with either vehicle (DMSO) or 4-hydroxytamoxifen (4-OHT) to generate F/F (Ctrl) and *Lpcat3*^IKO^ pBAs, respectively. qPCR analysis confirmed that *Lpcat3* expression was upregulated during the course of brown adipocyte differentiation (fig. S2A). The 4-OHT treatment regimen effectively depleted both LPCAT3 and AA-PE levels (Fig. 2, A and B). We observed a modest reduction in AA-PC species, yet their overall abundance was relatively minor (Fig. 2C). This lipidomic rewiring did not affect adipogenesis *per se* based on morphology and neutral lipid storage (fig. S2B). In addition, mtDNA copy number, OXPHOS subunits, and UCP1 protein levels were comparable between the genotypes (Fig. 2D and fig. S2, C and D).

**Fig. 2.**
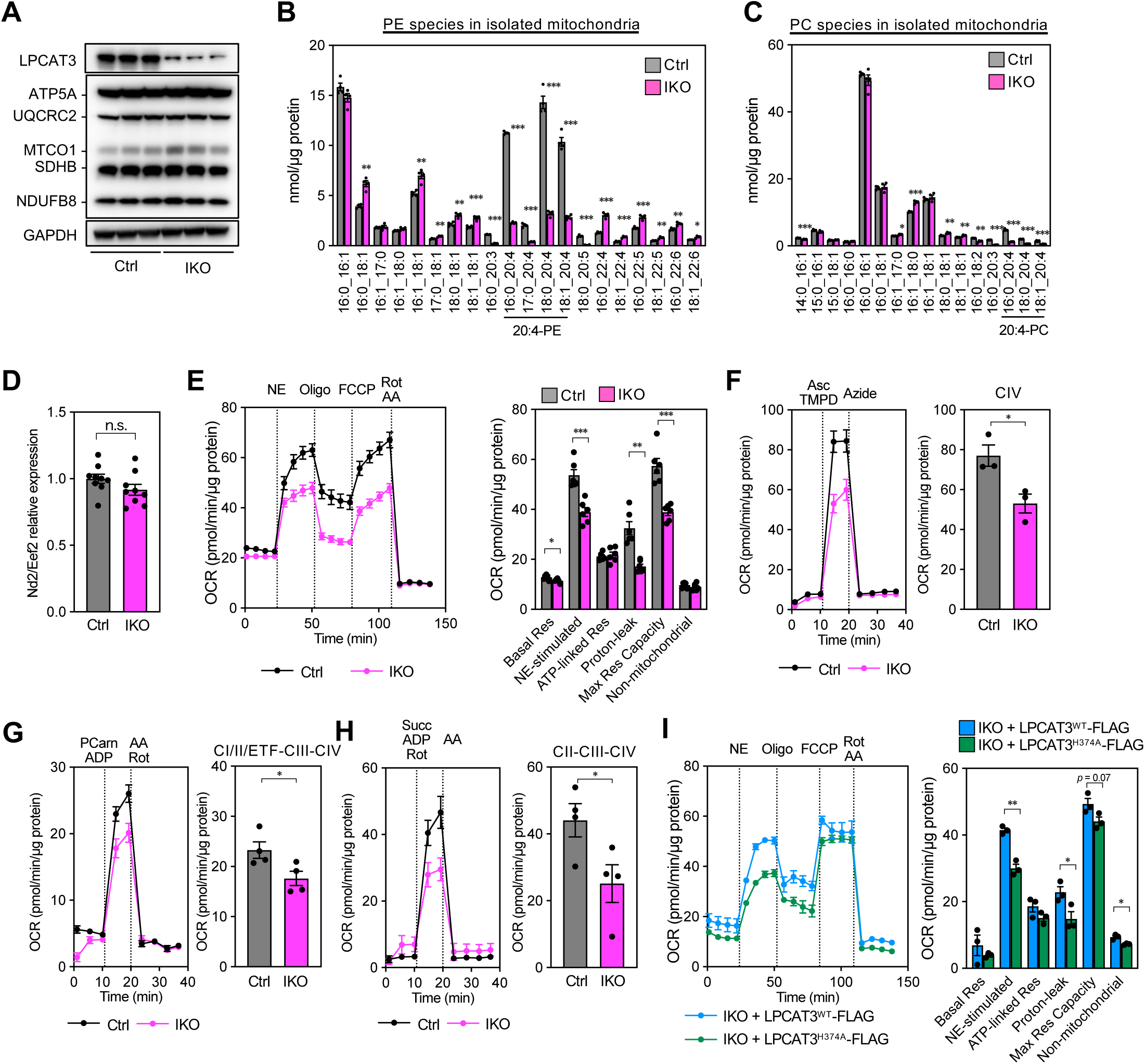
LPCAT3 catalytic activity is required for UCP1-dependent thermogenesis in brown adipocytes. (**A**) Western blot analysis of LPCAT3, UCP1, and OXPHOS subunits in *Lpcat3*^F/F^, *Cre*^ERT2^ pBAs treated with vehicle (Ctrl) or 4-OHT (*Lpcat3*^IKO^) and differentiated for 6 days (*n* = 3). GAPDH was used as a loading control. (**B, C**) Lipidomic analysis of (**B**) PE and (**C**) PC composition in Ctrl and *Lpcat3*^IKO^ pBA mitochondria (*n* = 4). (**D**) qPCR analysis of mtDNA copy number in Ctrl and *Lpcat3*^IKO^ pBAs (*n* = 9). (**E**-**H**) Seahorse respirometry performed on Ctrl and *Lpcat3*^IKO^ pBAs. (**E**) OCR traces (left) and quantification (right) after serial injections with NE (1 μM), oligomycin (OM, 4 μM), FCCP (10 μM), and rotenone/antimycin A (Rot/AA, 4/7.5 μM) (*n* = 6). (**F**) CIV-specific activity in NE-treated permeabilized Ctrl and *Lpcat3*^IKO^ pBAs, measured in the presence of antimycin A (4 μM) and GDP (1 mM), using ascorbate/TMPD as substrates (*n* = 3). OCR traces (left) and quantification (right) of CIV activity after injections with Asc/TMPD (700 μM/500 μM) and azide (33 mM). (**G**) CI/II/electron transfer flavoprotein (ETF)–CIII–CIV activity in permeabilized Ctrl and *Lpcat3*^IKO^ pBAs (*n* =4). OCR traces (left) and quantification (right) of CI/II/ETF–CIII–CIV respiration after serial injections with palmitoyl-carnitine/ADP (PCarn/ADP, 40 μM/4 mM) and Rot/AA (2/4 μM). (**H**) CII–CIII–CIV activity in permeabilized Ctrl and *Lpcat3*^IKO^ pBAs (*n* = 4). OCR traces (left) and quantification (right) of CII–CIII –CIV activity after serial injections with succinate/ADP/rotenone (Succ/ADP/Rot, 5 mM/4 mM/2 μM) and antimycin A (4 μM). (**I**) Seahorse respirometry performed on *Lpcat3*^IKO^ pBAs expressing Flag-tagged LPCAT3^WT^ or LPCAT3^H374A^ (*n* = 3). OCR traces (left) and quantification of respiratory states (right) after serial injections of NE (1 μM), OM (4 μM), FCCP (10 μM) and Rot/A, (4/7.5 μM). Data are presented as mean ± SEM. \**p*<0.05, \*\**p*<0.01, \*\*\**p*<0.001 by two-sided Welch’s *t*-test (B, C, D, E, F, G, H, I).

To mimic thermal stress *in vitro*, we treated Ctrl and *Lpcat3*^IKO^ pBAs with norepinephrine (NE)–an adrenergic receptor agonist. Under stimulated conditions, there were no gross changes in mitochondrial network morphology (fig. S2E). Moreover, we did not detect differences in the phosphorylation status of cAMP-dependent protein kinase A (PKA) substrates or lipolytic rate (fig. S2, F and G). However, despite comparable mitochondrial number and fuel mobilization, *Lpcat3*^IKO^ pBAs exhibited impaired NE-induced proton leak and maximal respiratory capacity relative to controls (Fig. 2E). We examined ETC capacity in detail by measuring the oxygen consumption rate (OCR) utilizing combinations of oxidative substrates and complex-specific inhibitors (fig. S2H). We found a consistent reduction in the OCR driven by OXPHOS complexes I/II-III-IV, II-III-IV, and IV in *Lpcat3*^IKO^ pBAs (Fig. 2, F to H), pointing to an integral defect in ETC efficiency.

To further strengthen the link between AA-PE levels and ETC activity, we reconstituted *Lpcat3*^IKO^ pBAs with Flag-tagged LPCAT3^WT^ or the catalytically inactive LPCAT3^H374A^ mutant (*41*). LPCAT3^WT^ re-expression in *Lpcat3*^IKO^ pBAs fully restored the membrane lipidome and AA-PE to levels detected in Ctrl pBAs (fig. S2I). Importantly, LPCAT3 enhanced NE-stimulated uncoupled respiration in a manner depending on its catalytic activity (Fig. 2I). This effect was not due to overt changes in pan-adipogenic or ETC indices between LPCAT3^WT^ and LPCAT3^H374A^-reconsituted *Lpcat3*^IKO^ pBAs (fig. S2, J to L). These data indicate that LPCAT3 activity bolsters UCP1-dependent uncoupled respiration and establish *Lpcat3*^F/F^, *Cre*^ERT2^ pBAs as a tractable *in vitro* model for studying the functional relevance of AA-PE in thermogenic capacity.

## Arachidonoyl-PE physically interacts with COX4I1 in brown adipocytes

Cold-induced lipidome remodeling may support ETC capacity by impacting membrane biophysical properties and/or specific lipid-protein arrangements (*32*). We utilized lipid-based chemical proteomics in combination with LPCAT3 loss-of-function strategies to address the thermoregulated (phospho)lipid-protein interactome (Fig. 3A). We chose an arachidonoyl lipid-based probe (AA-DA) that was readily incorporated into phospholipids (*42*). After UV irradiation, the crosslinked lipid-binding proteome was conjugated to a fluorescent reporter tag (Alexa 488-N_3_) by copper-catalyzed azide–alkyne cycloaddition (CuAAC or “click”). Analysis of probe targets by *in–gel* fluorescence scanning showed UV-dependent labeling that was sensitive to pre-treatment with either 4-OHT (*Lpcat3*^IKO^) or the LPCAT3 inhibitor (*R*)-HTS-3 (*43*), demonstrating that the AA-DA probe mimics the cognate acyl-CoA substrate of LPCAT3 (fig. S3, A and B). The signal intensity of a control oleic acid probe (OA-DA) was comparable between F/F and *Lpcat3*^IKO^ pBAs (fig. S3C).

**Fig. 3.**
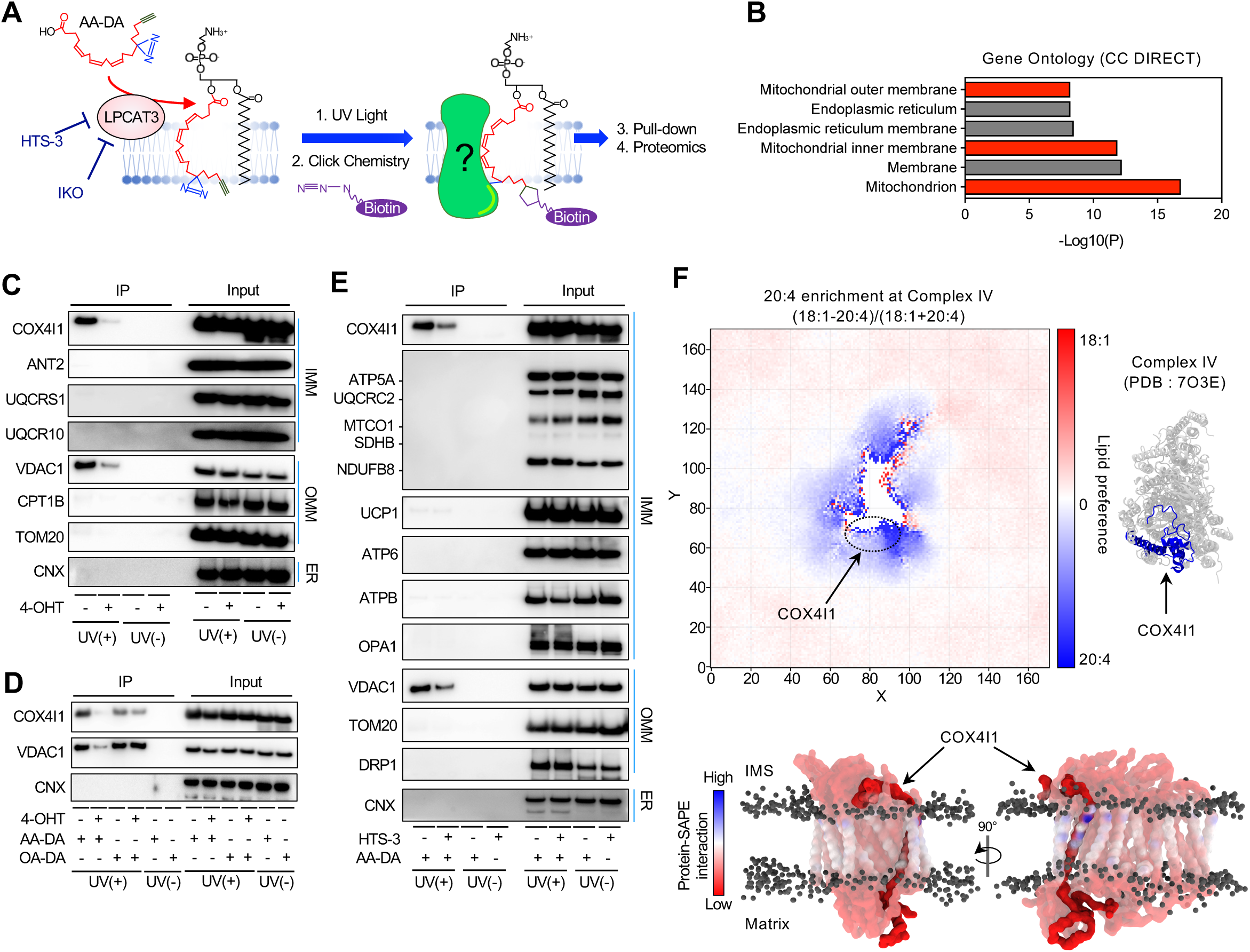
Arachidonoyl phospholipids selectively interact with COX4I1 in brown adipocytes. (**A**) Pipeline for the lipid-based chemical proteomics strategy for identifying candidates that physically interact with arachidonoyl phospholipids. *Lpcat3*^F/F^, *Cre*^ERT2^ (Ctrl) pBAs were incubated with the AA-DA probe (20 μM) for 1 h to generate AA-DA-containing phospholipids, followed by UV-irradiation to induce cross-linking. Membrane fractions were conjugated to either Alexa 488-N_3_ (*in–gel* visualization) or biotin (streptavidin pull-down) using click chemistry. As a negative control, *Lpcat3*^F/F^, *Cre*^ERT2^ pBAs were pre-treated with either 4-OHT (*Lpcat3*^IKO^) or the LPCAT3 inhibitor (*R*)-HTS-3 (10 µM). (**B**) Gene ontology analysis of 39 candidates identified in both conditions (Ctrl/IKO pBAs > 2 and DMSO/HTS-3-treated Ctrl pBAs > 2). (**C-E**) Validation of lipid–protein interactions in pBAs by immunoprecipitation (IP) and Western blot analysis. (**C**, **D**) Ctrl and *Lpcat3*^IKO^ pBAs were differentiated for 6 days, followed by incubation with AA-DA (20 µM) or OA-DA (20 µM). Proteins that were pulled-down by each respective photoaffinity lipid probe were detected by Western blotting. (**E**) Ctrl pBAs were differentiated for 6 days, followed by pre-treatment with either DMSO or (*R*)-HTS-3 (10 µM) for 2 h and incubation with AA-DA (20 µM). Proteins that were pulled-down by AA-DA were detected by Western blotting. Inputs are shown on the right and non-UV-irradiated (UV(-)) groups served as negative controls. (**F**) Top: 2D lipid preference analysis by coarse-grain molecular dynamics (CG–MD) simulations on a Complex IV crystal structure embedded in a virtual bilayer (comprised of 18:0/20:4-PE, 12.5%; 18:1/18:1-PE, 12.5%; 18:1/18:1-PC, 50%; tetra-18:2-CL, 25%). The area highly enriched for 20:4-over 18:1-containing phospholipids is shown in blue. Bottom: analysis of the COX4I1-PUFA interaction; contact frequency between 20:4 and Complex IV (amino acid residues interacting with PUFA chains highlighted in blue).

We next used click chemistry to map the interactome of LPCAT3-generated AA-DA phospholipids. Proteomics identified 39 AA-DA probe targets that were enriched in both Ctrl *vs*. *Lpcat3*^IKO^ pBAs and vehicle *vs*. HTS-3-treated pBAs (signal ratio ≧ 2.0; fig. S3D). Gene Ontology (GO) analyses revealed that the LPCAT3-dependent AA-DA interactome was enriched for IMM-localized proteins (Fig. 3B). In addition, 8 out of 39 proteins exhibited a preference for AA-DA over OA-DA chains (signal ratio ≧ 4/3; fig. S3D). Amongst these hits, COX4I1 was of particular interest given its role as a linchpin subunit of Complex IV (Cytochrome *c* oxidase, COX), the final and rate-limiting oxidase of the respiratory chain (*44, 45*). Immunoblot analysis validated that COX4I1 preferentially bound AA-DA over OA-DA, in an LPCAT3-dependent manner (Fig. 3, C to E). Only minor labeling was observed for abundant IMM and ETC components, illustrating the high specificity of the COX4I1 interaction. UCP1 exhibited weak AA-DA binding irrespective of LPCAT3 activity (Fig. 3E). This is in line with prior studies delineating free arachidonic acid as a potent allosteric UCP1 activator (*3*). The outer mitochondrial membrane (OMM)-residing VDAC1 interacted with both lipid probes (Fig. 3, C to E), possibly due to its moonlight activity as a lipid scramblase (*46, 47*).

Building on the observed dysregulated cytochrome *c* oxidase activity in *Lpcat3*^IKO^ pBAs, we further explored its interaction with arachidonoyl phospholipids. Coarse-grain molecular dynamics (CG–MD) simulations were performed on a Complex IV crystal structure (*48*) embedded within a virtual bilayer. A direct comparison of PE molecular species containing either arachidonoyl (SAPE) or oleoyl (DOPE) chains confirmed robust lateral partitioning of SAPE at the COX4I1 interface of Complex IV (Fig. 3F). This aligned with our proteomic data, highlighting preferential COX4I1 binding to AA-PE over oleoyl-PE. Such acyl chain selectivity was not detected for other OMM- and IMM-bound complexes (fig. S3, E to G). These data indicate that, instead of being ‘randomly’ distributed in the IMM, AA-PE engages in highly specific ETC interactions.

## Arachidonoyl-PE optimizes ETC efficiency during BAT thermogenesis

LPCAT3-derived AA-PE boosted UCP1-driven uncoupled respiration at the cellular level, leading us to explore its role in thermoregulatory physiology. Adipocyte-specific LPCAT3 deletion in mice did not impair their ability to sustain euthermia during acute cold challenge (fig. S4, A and B). Remarkably, however, when the spare capacity of BAT was first suppressed by pre-acclimating mice to TN (*49–51*), both *Lpcat3*^AKO^ and *Lpcat3*^BKO^ mice became hypothermic at a faster rate than their littermate controls (Fig. 4, A and B). Severe cold intolerance was also apparent in *Lpcat3^−/−^* female cohorts (Fig. 4, C and D), indicating that sex is not a major modifier of this phenotype. RNA-seq analysis of BAT before and after cold challenge revealed that the broad program of cold-inducible genes was largely unaffected by the loss of LPCAT3 (fig. S4, C and D). In accordance, downstream consequences of sympathetic BAT activation, including PKA signaling and adipose tissue lipolysis, were comparable between control and *Lpcat3*^BKO^ mice (fig. S4, E and F). These data suggest that β-adrenergic signaling remains intact in our LPCAT3 loss-of-function models.

**Fig. 4.**
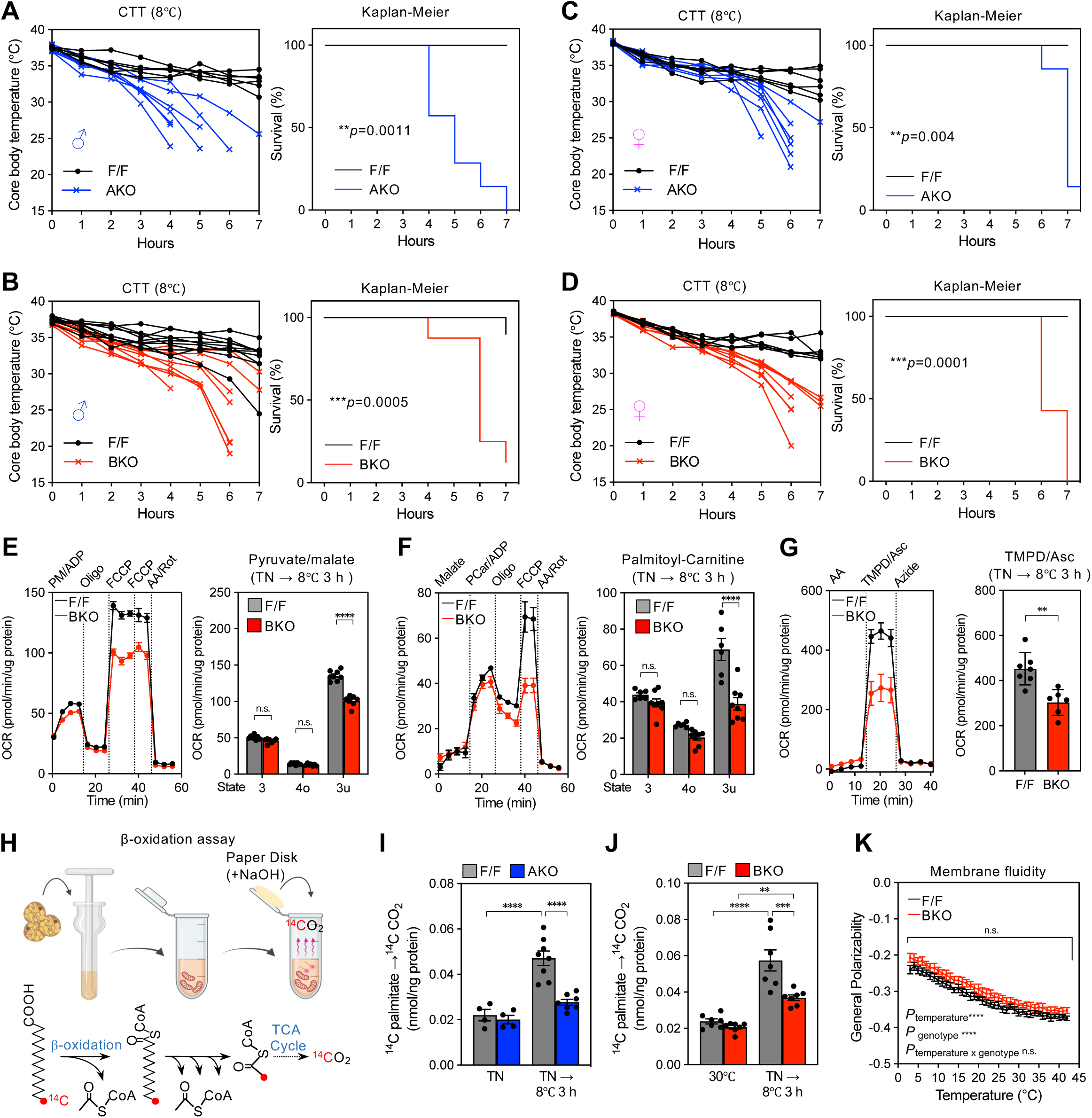
LPCAT3 deficiency impairs respiratory-dependent BAT thermogenesis during acute cold challenge. (**A-D**) Cold tolerance tests (CTTs) performed on 10-week-old (**A**) male F/F and *Lpcat3*^AKO^ mice (*n* = 7, 6), (**B**) male F/F and *Lpcat3*^BKO^ mice (*n* = 10, 8), (**C**) female F/F and *Lpcat3*^AKO^ mice (*n* = 6, 7), and (**D**) female F/F and *Lpcat3*^BKO^ mice (*n* = 6, 7). Mice were pre-acclimated to TN (30℃) for 2 weeks, followed by an acute CTT (8℃) and hourly monitoring of core body temperature (left). Core body temperatures ≤28°C were scored as events for the Kaplan-Meier survival curves (right). (**E-G**) Seahorse respirometry performed on isolated BAT mitochondria from F/F and *Lpcat3*^BKO^ mice acclimated to TN (30℃) and subjected to acute cold challenge (8℃) for 3 h (*n* = 6– 8/group). (**E**) CI/II–CIII–CIV activity measured in the presence of pyruvate (5 mM), malate (1 mM), ADP (4 mM), and GDP (1 mM). OCR traces (left) and quantification (right) of state 3 (ADP-stimulated), 4o (oligomycin leak), and 3u (FCCP-induced uncoupled/maximal) respiration after serial injections with oligomycin (Oligo, 4 μM), FCCP (4 μM), and rotenone/antimycin A (Rot/AA, 2/4 μM). (**F**) CI/II/ETF–CIII–CIV activity measured in the presence of malate (1 mM), GDP (1 mM), and using palmitoyl-carnitine as a substrate. OCR traces (left) and quantification (right) of respiratory (3/4o/3u) states after serial injections with palmitoyl-carnitine/ADP (PCarn/ADP, 40 μM /4 mM), Oligo (4 μM), FCCP (4 μM), and Rot/AA (2/4 μM). (**G**) CIV-specific activity measured in the presence of antimycin A (4 μM) and GDP (1 mM), using ascorbate/TMPD (Asc/TMPD) as substrates. OCR traces (left) and quantification (right) of CIV respiration after serial injections with Asc/TMPD (700 μM/500 μM) and sodium azide (33 mM). (**H-J**) *Ex vivo* fatty acid β-oxidation in crude BAT lysates, as assessed by the conversion of ^14^C-palmitate (PA) to ^14^CO_2_ (*n* = 4–8/group). (**H**) Graphic illustration of the *ex vivo* β-oxidation assay. BAT lysates were prepared from (**I**) F/F and *Lpcat3*^AKO^ or (**J**) F/F and *Lpcat3*^BKO^ mice kept at TN (30℃) or subjected to acute cold challenge (8℃) for 3 h. (**K**) Membrane fluidity of isolated mitochondria from cold-acclimated F/F and *Lpcat3*^BKO^ mice was assessed by C-Laurdan staining (*n* = 5/group). General polarizability (GP) was calculated at each 1°C increment from 3°C to 42°C. All animals described in this Figure were 10-week-old male mice and pre-acclimated to TN (30℃) for 2 weeks, unless stated otherwise. The figure was partially created using BioRender. Data are presented as mean ± SEM. \*\**p*<0.01, \*\*\**p*<0.001, \*\*\*\**p*<0.001 by two-sided Welch’s *t*-test (E, F, G), one-way ANOVA with Tukey’s multiple comparisons test (I, J), two-way ANOVA with Sidak’s multiple comparisons test (K), or Gehan-Breslow-Wilcoxon test (A, B, C, D).

To specifically narrow in on the AA-PE–COX interaction and its impact on respiratory-dependent thermogenesis, we conducted Seahorse extracellular flux analyses on isolated BAT mitochondria from F/F and *Lpcat3*^BKO^ mice following acute cold challenge. GDP was added to inhibit UCP1, allowing for the accurate analysis of electron transfer kinetics across varying respiratory states. When assayed with pyruvate/malate or palmitoyl-carnitine as oxidative substrates, *Lpcat3*^BKO^ BAT mitochondria exhibited diminished FCCP-induced maximal respiratory capacity relative to controls (Fig. 4, E and F). To probe COX-specific activity, we measured the OCR driven by TMPD/ascorbate in the presence of antimycin A (Complex III inhibitor). Indeed, electron flux through Complex IV was greatly reduced in *Lpcat3*^BKO^ BAT mitochondria (Fig. 4G). As a complementary approach, we evaluated the β-oxidation capacity of BAT *ex vivo* by monitoring the conversion of ^14^C-palmitic acid to ^14^CO_2_ (Fig. 4H). Consistent with our bioenergetic analyses, *Lpcat3^−/−^* BAT showed a diminished burst of cold-stimulated β-oxidation (Fig. 4, I and J). Importantly, these effects appeared to be independent of gross changes in membrane fluidity, as determined by C-Laurdan staining across a range of temperatures (Fig. 4K). Moreover, the defect in ETC capacity was cold-specific and arose in the absence of variations in key mitochondrial indices, including ETC content and assembly, cristae morphology, and UCP1-mediated proton currents (fig. S4, G to N). These data suggest that the AA-PE–COX interaction facilitates a catalytic state conducive to efficient electron transfer during respiratory-dependent BAT thermogenesis.

## Arachidonoyl-PE regulates ETC form and function in thermogenic fat

Lastly, we assessed the consequence of impaired electron transfer capacity on BAT physiology and mitochondrial bioenergetics under cold adaptation. *Lpcat3*^BKO^ mice housed at 23 ℃ exhibited typical BAT morphology and ultrastructure relative to F/F controls (fig. S5, A to C). Accordingly, there were no major differences in the thermogenic gene program, UCP1 protein levels, OXPHOS abundance, or substrate utilization (fig. S5, D to H). Conversely, cold-acclimated *Lpcat3*^BKO^ mice had larger and paler BAT relative to their floxed littermates (Fig. 5A). Histological examination confirmed a more unilocular appearance of *Lpcat3^−/−^*BAT depots (Fig. 5B). We also observed overt signs of dysmorphic brown fat mitochondria, as evidenced by irregular cristae density and/or vesiculation (Fig. 5C). Despite abnormal BAT morphology, the thermogenic gene program and iWAT “browning” responses were largely intact in *Lpcat3*^BKO^ mice (fig. S6, A to C).

**Fig. 5.**
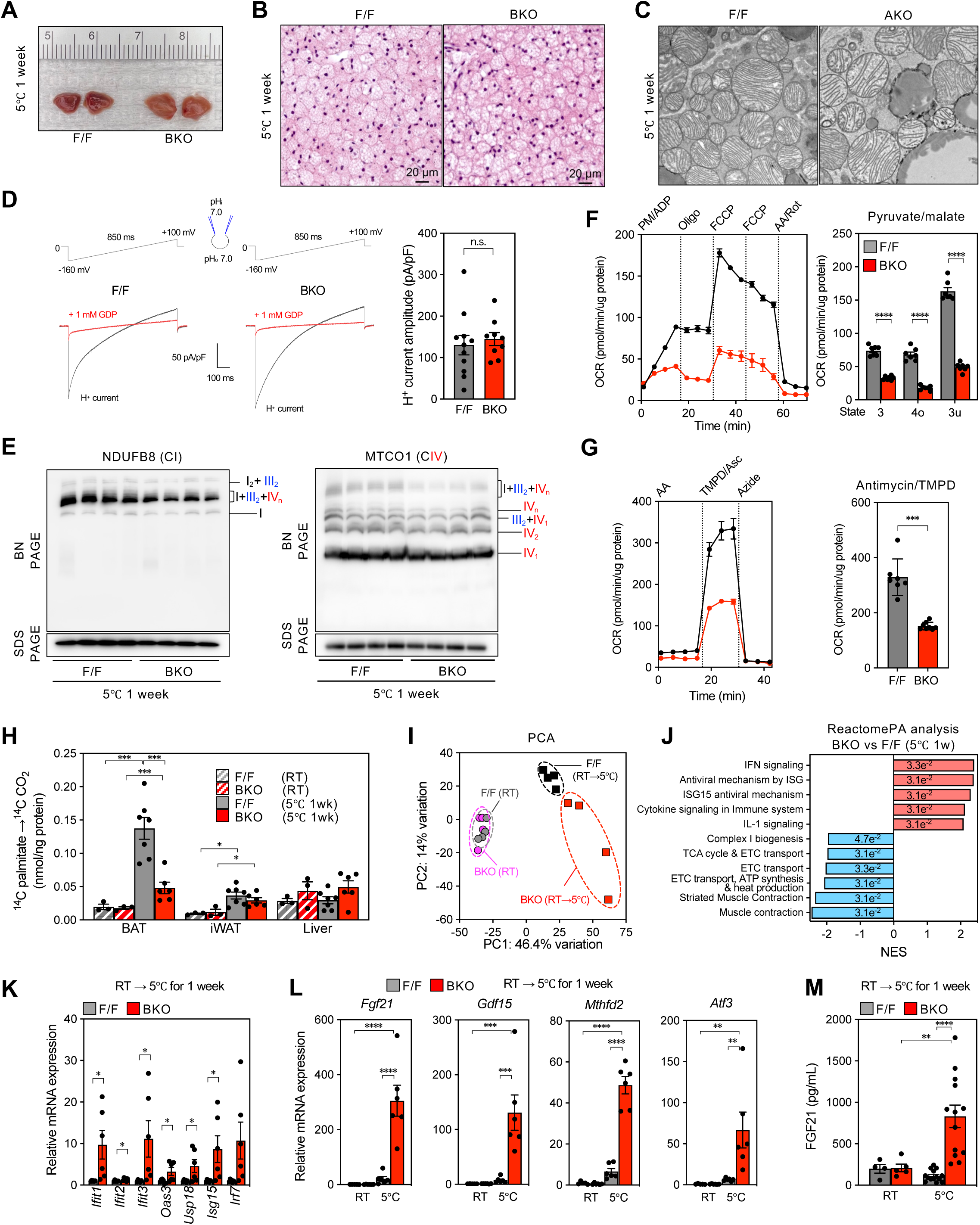
LPCAT3 deficiency in BAT triggers defects in ETC form and function upon cold adaptation. (**A-C**) Gross appearance (**A**), H&E staining sections (**B**), and electron micrographs (**C**) of BAT from 10-week-old male F/F and *Lpcat3*^BKO^ mice acclimated to cold exposure (5℃) for 7 days. (**D**) Electrophysiological recording of proton (H^+^) currents across the IMM of BAT-derived mitoplasts from cold-acclimated F/F and *Lpcat3*^BKO^ mice. Top: voltage ramp protocol. Bottom: representative traces showing proton currents before (black) and after (red) addition of 1 mM GDP. Right: summary of H^+^ current amplitude (*n* = 9-11/group). (**E**) BN–PAGE Western blot analysis of isolated BAT mitochondria from F/F and *Lpcat3*^BKO^ mice acclimated to 5℃ for 7 days to evaluate SC assembly using OXPHOS complex-specific antibodies: CI (NDUFB8) or CIV (MTCO1). SDS–PAGE Western blot analysis was performed on samples akin to those for BN-PAGE to serve as loading controls (*n* = 4/group). (**F, G**) Seahorse respirometry performed on isolated BAT mitochondria from F/F and *Lpcat3*^BKO^ mice acclimated to 5℃ for 7 days (*n* = 7/group). (**F**) CI/II–CIII–CIV activity was measured in the presence of pyruvate (5 mM), malate (1 mM), ADP (4 mM), and GDP (1 mM). OCR traces (left) and quantification (right) of the respiratory (3/4o/3u) states after serial injections with Oligo (4 μM), FCCP (4 μM), and Rot/AA (2/4 μM). (**G**) CIV-specific activity measured in the presence of antimycin A (4 μM) and GDP (1 mM), using Asc/TMPD as substrates. OCR traces (left) and quantification (right) of CIV respiration after serial injections with Asc/TMPD (700 μM/500 μM) and azide (33 mM). (**H**) *Ex vivo* fatty acid β-oxidation in crude BAT, iWAT and liver lysates, as assessed by the conversion of ^14^C-PA to ^14^CO_2_ (*n* = 4–8/group). Crude tissue lysates were prepared from F/F and *Lpcat3*^BKO^ mice either housed at 23℃ or acclimated to 5℃ for 7 days. (**I**, **J**) RNA-seq was performed on BAT from F/F and *Lpcat3*^BKO^ mice either kept at 23℃ or adapted to 5℃ for 7 days (*n* = 4/group). (**I**) PCA of the BAT transcriptome. (**J**) ReactomePA analysis of the up- and down-regulated pathways (normalized enrichment score, NES). (**K**) qPCR analysis of IFN-related transcript levels in BAT from F/F and *Lpcat3*^BKO^ mice acclimated to 5℃ for 7 days (*n* = 6, 6). (**L**) qPCR analysis of ISR^mt^-related transcript levels in BAT from F/F and *Lpcat3*^BKO^ mice kept at 23℃ or acclimated to 5℃ for 7 days (*n* = 6/group). (**M**) Plasma FGF21 levels in F/F and *Lpcat3*^BKO^ mice kept at 23℃ or acclimated to 5℃ for 7 days (*n* = 4–14/group). All animals described in this Figure were 10-week-old males and reared at 23℃. Data are presented as mean ± SEM. \**p*<0.05, \*\**p*<0.01, \*\*\**p*<0.001, \*\*\*\**p*<0.0001 by two-sided Welch’s *t*-test (D, F, G, K), or one-way ANOVA with Tukey’s multiple comparisons test (H, L, M).

The selective accrual of AA-PE species in BAT mitochondria during cold exposure prompted to re-evaluate UCP1 function. Under cold acclimation, BAT-derived mitoplasts from control and *Lpcat3*^BKO^ mice exhibited comparable UCP1-driven proton currents (Fig. 5D). Accordingly, LPCAT3 loss did not affect UCP1 protein levels and OXPHOS subunits (fig. S6D). However, blue native–polyacrylamide gel electrophoresis (BN–PAGE) analysis revealed a specific assembly defect of the I/III_2_/IV_n_ holo-complex, known as ‘respirasome’ (*52–54*), in BAT of cold-adapted *Lpcat3*^BKO^ mice (Fig. 5E and fig. S6E). This finding was cross-validated using complex-specific antibodies and *in–gel* activity assays. Complex II existed as a ‘free-floating’ monomer and its distribution was comparable between the genotypes (fig. S6F). Decreased respirasome levels in *Lpcat3*^BKO^ mitochondria were associated with severely impaired electron transfer capacity (Fig. 5, F to H).

Besides conferring a kinetic advantage, SCs help curtail reactive oxygen species (ROS) formation in the respiratory chain (*55, 56*). To assess the global molecular consequence of impaired SC assembly and ETC function, we performed RNA-seq analysis of BAT from cold-adapted *Lpcat3*^BKO^ and control mice. Principal component analysis (PCA) revealed a cold-induced shift in the transcriptomes (Fig. 5, I and J), partly due to reduced expression of mitochondrially-encoded ETC subunits in *Lpcat3^−/−^* BAT (fig. S6, G and H). Conversely, genes linked to interferon ψ (IFNψ) signaling and the mitochondrial integrated stress response (ISR_mt_) were coordinately upregulated in *Lpcat3^−/−^*BAT (Fig. 5, J to L). Indeed, qPCR analysis showed that transcripts of ISR_mt_-related genes (*e.g*., *Fgf21*, *Gdf15*, *Mthfd2*, *Atf3*) increased over time in a cold-inducible manner (Fig. 5L and fig. S6I). Accordingly, BAT-derived FGF21 was robustly elevated in the plasma of cold-adapted *Lpcat3*^BKO^ mice (Fig. 5M and fig. S6J). ISR_mt_ activation in *Lpcat3^−/−^* BAT was reinforced by increased ATF4 protein expression and augmented eIF2α phosphorylation (fig. S6K). These data suggest that AA-PE accrual in brown fat mitochondria optimizes ETC form and function in a thermal stress-dependent manner.

## Discussion

Cold-induced remodeling of the BAT lipidome has been recognized for decades (*16–18*). This unique trait of thermogenic adipocytes is conserved across animal species, yet its exact function in thermoregulatory physiology has been a longstanding mystery. Does the cold-induced shift in acyl chain profiles support ETC capacity through influencing membrane biophysical properties and/or by engaging specific (phospho)lipid–protein interactions? Here, we identify LPCAT3 as the remodeling enzyme required for the thermal stress-dependent accrual of AA-PE in brown fat mitochondria. Lipid-based proteomics, CG–MD simulations, and bioenergetic analyses revealed that AA-PE partitions at the COX4I1 interface of Complex IV, enhancing its catalytic efficiency. The near-complete loss of AA-PE in *Lpcat3*^BKO^ mice impairs respiratory-dependent BAT thermogenesis and predisposes to cold-induced hypothermia. Thus, our results reveal a cold-regulated lipid–protein interaction as a gating factor in UCP1-mediated uncoupled respiration. The ubiquitous expression of COX4I1 raises the prospect that the AA-PE–COX interaction might be leveraged in mitochondrial pathologies, such as rare monogenic disorders, obesity, and ageing.

Prior work from our lab and others has revealed tissue-specific roles of LPCAT3 in key lipid utilization pathways. We have shown that LPCAT3 activity supports triglyceride storage in unilocular white adipocytes through enhancing lipid droplet (LD) budding size (*40*). Paradoxically, cold adaptation was associated with *increased* LD accrual in *Lpcat3^−/−^* BAT. This apparent discrepancy may be explained by the emergence of BAT-specific mechanism(s) (*e.g.*, CLSTN3β) that uncouple LPCAT3 activity from LD size regulation (*49, 57*). In addition, compromised ETC efficiency and thermogenic β-oxidation in cold-adapted *Lpcat3^−/−^* BAT would favor LD accrual regardless of a defect in budding size. A similar overt steatotic phenotype has been observed in *Lpcat3^−/−^* livers (*38, 39*), arising from impaired VLDL assembly and triglyceride export. Of note, a recent study by Ye *et al*. (*58*) has sought to reconcile hepatic LPCAT3 activity with mitochondrial function in a murine NASH model. In contrast to our study, they reported that LPCAT3 loss-of-function was associated with mitochondrial hyperactivity and ROS formation. The basis of this discrepancy is unclear, but it is conceivable that the effects Ye *et al*. observed are secondary to exaggerated steatosis and lipotoxicity in *Lpcat3^−/−^*livers. Ye *et al*. (*58*) did not specifically address the mechanisms underlying ETC dysfunction and/or the contribution of distinct phospholipid species (*e.g*., AA-PE).

A central dogma in the field is that cold-induced mitochondrial lipidome remodeling supports BAT thermogenesis by ensuring membrane fluidity and/or engaging specific lipid-protein arrangements (*25*). Indeed, the biophysical properties of membranes are influenced by the length and unsaturation degree of its acyl chains (*33*). To our surprise, LPCAT3 loss did not impact general fluidity of BAT mitochondrial membranes, though more localized changes may still occur. In accordance, we have shown that AA-PE selectively partitions at the COX4I1 interface of Complex IV, the final and rate-limiting oxidase of the respiratory chain. COX4I1 is a single-pass transmembrane (TM) protein that interacts with the catalytic subunit MT-CO1. COX4I1 also facilitates the docking of Complex III-derived CYT*C* to MT-CO2 (*59*), thus finetuning COX activity, and by extension, ETC flux. Additionally, COX4I1 is phosphorylated at Ser58 in a PKA-dependent manner (*60*). This modification derepresses COX by preventing its allosteric inhibition by ATP, further establishing COX4I1 as a key regulatory hub for ETC capacity. Of note, the TM domain of COX4I1 bears homology to that of yeast MGA2, which acts as a lipid-packing sensor in the ER membrane (*61*). Thus, the AA-PE–COX4I1 interaction may locally preserve membrane fluidity, safeguarding COX activity and ETC flux under cold stress.

LPCAT3 activity regulates ETC form and function in a thermal stress-dependent manner. It appears that LPCAT3 becomes essential only once a certain threshold of ETC activity is reached. ETC capacity is known to vary across metabolic tissues, with BAT showing the greatest OCR per unit of mitochondria (*62*). The cold-regulated AA-PE–COX4I1 interaction likely underpins the beneficial impact of LPCAT3 activity on BAT thermogenesis. Supporting this, prior work has shown that PE partitions at the interface of a crystalline bovine heart Complex IV dimer (*63*), underlining its role in COX assembly and function. Furthermore, COX activity directly correlates with mitochondrial PE content in PE *N*-methyltransferase (*Pemt*)-null hepatocytes (*64*). Of interest, PE has recently been proposed as a temperature-responsive rheostat controlling UCP1 activity (*28*). However, this is solely based on studies using BAT-specific PS decarboxylase (*Psd*)-KO mice, which exhibit general deficits in mitochondrial PE levels, mtDNA copy number, cristae density, and thermogenic gene profiles. Our data suggest that the cold-inducible shift in PE composition is not a determinant of UCP1 function *per se*, but rather supports BAT thermogenesis by engaging specific interactions with respiratory complexes. Our results align with a study by Shin *et al*. (*32*) who delineated cold-inducible conformational switches in brown fat respiratory SCs (CI:III_2_) to electron transfer-favorable catalytic states. LPCAT3-derived AA-PE localizes to the COX4I1 subunit of Complex IV, which is juxtaposed to CIII within SCs (*65–70*). Our data point to a targeted effect on COX activity; however, contributions to broader SC dynamics beyond the resolution of BN–PAGE cannot be ruled out.

In mammals, PE formation occurs through the ER-resident *de novo* CDP-ethanolamine pathway or the IMM-resident PS decarboxylation pathway. Based on stable isotope studies (*71*), mitochondrial PE appears to be generated exclusively through the conversion of PS to PE by PSD. Of interest, prior work has suggested LPCAT3 has lyso-PS acyltransferase activity with 20:4-CoA (*41*). Therefore, LPCAT3 may catalyze the formation of AA-PS in mitochondria-associated membranes (MAMs), followed by transport to the IMM and subsequent conversion to AA-PE. PSD preferentially synthesizes PUFA-PE species (*e.g.*, 18:0_20:4, 18:0_22:6), while the *de novo* pathway forms PE with mono- or di-unsaturated fatty acyl chains (*e.g.*, 16:0_18:2, 18:1_18:2) (*72*). This could account for the selective accrual of AA-PE over AA-PC in brown fat mitochondria. On the other hand, ER-derived AA-PE may be transported directly to the IMM. Although phospholipid scramblases are known to localize to MAMs, an enzyme capable of moving PE across the OMM has yet to be established. Intriguingly, several OMM insertases (*e.g.*, Tom40, MTCH1/2, Sam50) and the anion-channel VDAC1 are thought to possess inherent lipid scrambling activity (*46, 47*). Based on our lipid-based proteomics data, it is tempting to speculate that VDAC1 may act as a PE scramblase in mammals.

Mitochondria in eukaryotic cells appear to have originated from engulfed proteobacteria. In support of this, CL and PE are the dominant phospholipid classes in mitochondria (*73*). Bacteria have long been known to incorporate unsaturated fatty acids into their membrane lipidome, thereby maintaining fluidity and ensuring survival in cold temperatures (*74*). Thermal stress-dependent phospholipid remodeling and IMM unsaturation in brown adipocytes may thus reflect evolutionary vestiges from when mitochondria were independent bacteria.

## Acknowledgments

We thank current and former members of the Tontonoz, Tarling-Vallim, Edwards, Villanueva, and Bensinger labs for valuable discussions and for sharing reagents. RNA-seq analyses were performed at the Technology Center for Genomics & Bioinformatics at UCLA.

## Funding

Japan Society for the Promotion of Science abroad (YS)

Osamu Hayaishi Memorial Scholarship for Study Abroad (YS)

American Diabetes Association (1-19-PDF-039 to M.J.T.)

National Institutes of Health (R01 DK129276 and DK136150 to P.T.; HL139725 to S.G.Y.; R35GM143097 to A.M.B.; GM142960 to I.B.)

Swiss National Science Foundation (grants 310030_219264 to S.V.)

European Research Council under European Union’s Horizon 2020 research and innovation program (grant agreement no. 803952, to S.V.)

Pew Scholars in Biomedical Sciences (25PRE1374054 to A.M.B.)

PID2021-127278NB-I00, funded by MCIN/AEI/ 10.13039/501100011033 and by “ERDF A way of making Europe”, by the European Union, to M.L.

## Author contributions

Conceptualization: YS, MJT, PT

Methodology: YS, MJT, CRR, LC, AB, AHB, BM, KJW, LS, MS, TAW, WC, JPW, IB, AMB, BFC, SMH, CA, DAF, ML, SV

Investigation: YS, MJT, CRR, MJJ, LC, AB, AHB, MGM, BM, BMS, AW, KJW, LS, MS, TAW, WC, AMB

Visualization: YS, MJT, SV, PT

Funding acquisition: YS, MJT, SV, SGY, PT

Project administration: PT

Supervision: ML, OSS, SV, PT

Writing – original draft: YS, MJT, PT

Writing – review & editing: YS, MJT, PT

## Data and materials availability

Source data for all figures are provided with the paper. Sequencing data have been deposited to GEO (GSE294663). All unique biological materials used are readily available from the authors or from standard commercial sources.

## Supplementary Materials

Materials and Methods

Figs. S1 to S6

## Materials and Methods

### Animal models and diets

*Lpcat3*^AKO^ and *Lpcat3*^BKO^ models were generated as described (*40*). The *Lpcat3*^IKO^ model was generated by breeding *Lpcat3*^F/F^ mice to *Rosa26*-CreERT2 mice (Jackson Labs, B6.129-*Gt(ROSA)26Sor^tm(cre/ERT2)Ty^j/*J, Stock 008463). Animal studies were performed in accordance with ethical regulations and protocols approved by the UCLA Institutional Animal Care and Use Committee (IACUC). Unless stated otherwise, all animal studies used age-matched littermates on a C57BL/6J background (Jackson Labs) group-housed at 3-5 mice per cage in a climate-controlled, specific-pathogen-free animal facility at 23°C with a 12:12-h light-dark cycle (lights on; 6 AM) and free access to chow diet (LabDiet, 5001) and water.

### Cold tolerance test

Cold tolerance tests were typically performed on age matched 9–11-week-old littermates. Thermoneutrality and cold studies were performed in climate-controlled incubators (Thermo Scientific) with a 12:12 h light-dark cycle (lights on; 6 AM) kept at either 28-29°C or 5-10°C, respectively. For the acute cold tolerance studies, mice were pre-acclimated to 28-29°C for 2 weeks. Mice were then fasted for 2 h (8 to 10 AM) at TN and singly-housed in pre-chilled cages with 25 g wood chip bedding, 1x cotton nesting, and *ad libitum* access to water. Mice were placed at 8°C for 7 h, during which core body temperature was measured hourly using the BAT-12 microprobe thermometer (Physitemp). Mice were euthanized once their core body temperature reached <28°C. The cold acclimation protocol was as follows: mice reared at room temperature (23°C) were singly-housed in cages with 50 g wood chip bedding, 2x cotton nesting, and *ad libitum* access to water and normal chow diet at 5°C for 7 days.

### Tissue and plasma harvesting

Mice were euthanized by cervical dislocation secondary to isoflurane overdose after 2 h of fasting. Blood was collected by cardiac puncture in BD microtainer K2-EDTA collection tubes (Fisher Scientific). Plasma was collected by centrifugation at 2,000 × *g* for 10 min at 4°C and stored at –80°C. Organs were quickly dissected and stored in formalin or snap-frozen in liquid nitrogen and stored at –80°C until further analysis. Plasma TG, NEFA, glycerol, and FGF21 levels were measured with Wako L-Type TG M Kit (Fujifilm), Wako HR Series NEFA-HR Kit (Fujifilm), Free Glycerol Reagent (F6428, Sigma) and Mouse/Rat FGF21 Quantikine ELISA Kit (MF2100, R&D Systems) according to the manufacturer’s instructions.

### Histological analysis

The harvested tissues were submerged in phosphate-buffered 4% formalin (Fisher Scientific) in cassettes for 1 week. Tissue cassettes were rinsed with running tap water for 15 min and stored in 70% ethanol until further processing. The paraffin embedding followed by tissue sectioning (5 µm) and H&E staining were done by the UCLA Translational Pathology Core Laboratory. Images were taken with a Zeiss Axioskop 2 Plus microscope.

### Western blot analysis

Whole-tissue and whole-cell lysates were made using RIPA buffer (Boston BioProducts) supplemented with protease and phosphatase inhibitor cocktails (Roche). Protein concentrations were determined by BCA Protein Assay Kit (Pierce). Equal amounts of protein were loaded onto NuPAGE 4-12% Bis-Tris gels (Invitrogen) and transferred to Amersham Hybond P 0.45 µm PVDF membranes (Cytiva). Membranes were blocked with 5% milk or 5% BSA in TBS-T (0.05% Tween-20) for 1 h at RT. Primary antibodies were diluted in 5% BSA in TBS-T and incubated with membranes overnight at 4°C under gentle agitation. The next day, membranes were washed three times with TBS-T at RT, 10 min per wash. Secondary antibodies were diluted in 5% milk (1:8,000) in TBS-T and incubated with membranes for 1 h at RT. After three TBS-T washes, the membranes were incubated with Immobilon Forte Western HRP substrate (Millipore) for 1-3 min. Imaging was performed by ChemiDoc system (BIO-RAD).

Antibodies used in this study were the following: Phosphor-PKA substrate (RRXS*/T*) (100G7E; #9624), COX4 (#4844), phospho-eIF2α (Ser51) (119A11; #3597), EIF2α (D7D3; #5324), ATF4 (D4D8; #11815), VDAC (D73D12; #4661), phospho-HSL (Ser660) (#45804), HSL (#4107), ANT2 (E2B9D; #14671), DRP1 (4E11B11; 14647) were from Cell Signaling Technology; TOM20 (#11802-1-AP), ATPB1 (#17247-1-AP), ATP6 (#55313-1-AP), CPT1B (#22170-1-AP) were from ProteinTech; OXPHOS rodent antibody cocktail (ab110413), CNX (ab10286), MTCO1 (#1D6E1A8; ab14705), UQCRSF1 (#5A5; ab14746), UCP1 (ab10983) were from Abcam; UQCR10 (NBP2-93778) were from Novus Biologicals; NDUFB (20E9DH10C12; 459210) was from Invitrogen; OPA1 (BDB612606) was from BD Biosciences. The polyclonal antibody for LPCAT3 was generated as described (*40*).

### Fatty acid β-oxidation assay

*Ex vivo* β-oxidation were performed as described with minor modifications (*75*). Briefly, excised tissues were homogenized using a motorized pestle (800 rpm, 10 strokes) in ice-cold STE buffer (0.25 M sucrose, 1 mM EDTA, and 10 mM Tris-HCl, pH 7.4). Homogenates (BAT (one lobe/0.5 ml STE), iWAT (one lobe/1 ml STE), or liver (200 mg/0.5 ml STE)) were decanted into 15 mL tubes, followed by centrifugation at 420 × *g* for 10 min at 4°C. A portion of the supernatant was collected for protein determination (BCA protein assay kit, Pierce). 30 μl supernatant was transferred to new Eppendorf 1.5 mL tubes (duplicate samples), followed by addition of 370 μl oxidation assay buffer (100 mM sucrose, 10 mM Tris-HCl pH 7.4, 5 mM KH_2_PO_4_, 0.2 mM EDTA pH 8.0, 80 mM KCl, 1 mM MgCl_2_, 2 mM L-carnitine, 0.1 mM malate, 0.05 mM coenzyme A, 2 mM ATP, 1 mM DTT, and 0.7% essentially fatty acid-free BSA) containing 0.5 μCi [^14^C]-palmitic acid (Perkin Elmer). After the reaction at 37°C for 30 min, the whole reaction mixture was transferred to new Eppendorf 1.5 mL tubes containing 200 μl of 6 M HCl and a paper disc pre-wetted with 20 μl of 1 M NaOH in the lid for ^14^CO_2_ trap. Tubes were closed and incubated for 1 h at RT. The radioactivity of ^14^CO_2_ on the paper disc was quantified by a scintillation counting. Counts were normalized for protein content.

### Pure mitochondria isolation for lipidomics analysis

Pure mitochondria isolation from BAT was performed as previously described with some modifications (*76*). Freshly harvested iBAT from 3 mice were placed in ice-cold MES buffer (225 mM mannitol, 75 mM sucrose, 30 mM Tris-HCl (pH 7.4), 0.5 mM EGTA, 2% essentially fatty acid-free BSA) and washed with SMTE buffer (225 mM mannitol, 75 mM sucrose, 30 mM Tris-HCl (pH 7.4), 0.5 mM EGTA, 0.5% essentially fatty acid-free BSA) three times. Fat pads were minced using scissors and homogenized using a motorized pestle (800 rpm, 20 strokes) in ice-cold MES buffer (2.5 ml/sample). Homogenates were decanted into 2 mL tubes, followed by centrifugation at 8500 × *g* for 10 min at 4°C. The supernatant was decanted rapidly, and the sides of the tube were wiped to remove lipids. The pellet was resuspended with 2 ml of MES buffer, followed by centrifugation at 740 × *g* for 5 min at 4°C. The supernatant (mitochondria fraction A) was transferred to a clean tube and kept on ice. The remaining pellet was resuspended in 2 ml of MES buffer, followed by centrifugation at 740 × *g* for 5 min at 4°C. After collecting supernatant (mitochondria fraction B), repeat this step once more to collect supernatant (mitochondria fraction C). Mitochondria fractions A-C were spun down at 8500 × *g* for 10 min at 4°C and the pellet was pooled by resuspending with 1 ml fresh MES buffer. These crude mitochondria suspensions were spun down at 10000 × *g* for 10 min at 4°C, and the pellet was resuspended in fresh SMTE buffer. These wash steps were repeated 4 times to remove all traces of microsomal contamination, and the remaining mitochondrial pellet was resuspended in 0.5 ml ice-cold MRB buffer (250 mM mannitol, 5 mM HEPES (pH 7.4), 0.5 mM EGTA). The Percoll density gradient was prepared by adding 3.2 ml Percoll medium (225 mM mannitol, 25 mM HEPES (pH 7.4), 1 mM EGTA, 30% Percoll (vol/vol)) to a 5-ml thin-wall polyallomer ultracentrifuge tube (#344057, Beckman), layered with 0.5 ml crude BAT mitochondria, and finally layered with 1ml MRB buffer. After centrifugation (95000 × *g* for 30 min at 4°C), the mitochondrial band near the bottom of the Percoll density gradient was collected and washed with 10 times volume of MRB buffer by 7500 × *g* for 10 min. The obtained mitochondrial pellet was resuspended in PBS, snap-frozen, and stored at –80°C until further analysis.

### Lipidomic analysis

#### Shotgun lipidomics (Fig. 1 A-D, Fig. 2 B, C, fig. S2 I)

Frozen tissues were weighed for normalization and placed in 2 mL homogenizer tubes with 6 ceramic beads (2.8mm; Omni #19-628) and 750 μl PBS. Tissues were homogenized in Omni Bead Ruptor Elite (3 cycles of 10 sec at 5 m/s with a 10 sec dwell-time). Homogenates (1-10mg) were used for lipid extraction with a modified Bligh and Dyer method (*77*). An internal standard mixture consisting of 75 lipid standards across 17 subclasses based on Ultimate Splash ONE was added to each sample prior to the lipid extraction (Avanti 330820, Avanti 861809, Avanti 330729, Avanti 330727, Avanti 791642, Avanti 330726). Following two successive extractions, pooled organic phases were dried down in a Thermo SpeedVac SPD300DDA using ramp setting 4 at 35°C for 45 min for 90 min. Lipid samples were resolved in methanol/dichloromethane (1:1) with 10 mM ammonium acetate and transferred to vials (Thermo 10800107). For TG-rich samples, a neutral lipid removal was performed to enable sensitive polar lipid detection. In brief, the dried down whole lipid extracts were resuspended in methanol/hexane (1:1), mixed, spun down, and the top hexane layer was discarded. Remaining lower layer was dried up and resuspended as described above. The samples were analyzed using the Sciex 6500+ equipped with Differential Mobility Device (DMS) device with an expanded targeted acquisition list consisting of 1450 lipid species across 17 subclasses (or an original acquisition list of 1100 lipids across 13 subclasses). DMS on Lipidyzer was calibrated with EquiSPLASH LIPIDOMIX (Avanti 330731). Data analysis was performed on an in-house analysis platform similar to the Lipidyzer Workflow Manager (*78*).

#### LC-MS/MS-based lipidomics (Fig. 1 I, J, fig. S1 F-I)

BAT and mitochondria were subjected to a modified Bligh-Dyer lipid extraction in the presence of lipid class internal standards including 1,2-diarachidoyl-sn-glycero-3-phosphocholine, 1,2-dimyristoyl-sn-glycero-3-phosphoethanolamine, 1,2-dimyristoyl-sn-glycero-3-phosphoserine, and 1’,3’-bis[1,2 dimyristoyl-sn-glycero-3-phospho]-glycerol (*77, 79*). For phosphatidylcholines and phosphatidylserines, lipid extracts were diluted in methanol/chloroform (4/1, v/v) and molecular species were quantified using electrospray ionization mass spectrometry on a triple quadrupole instrument employing shotgun lipidomics methodologies.

Phosphatidylcholine molecular species were quantified as chlorinated adducts in the negative ion mode using neutral loss scanning for 50 amu (collision energy = 24eV). Phosphatidylserine molecular species were quantified in the negative ion mode using neutral loss scanning for 87 amu (collision energy = 25eV).

Phosphatidylethanolamine molecular species were first converted to their fMOC derivatives and then were quantified in the negative ion mode using neutral loss scanning for 222.2 amu (collision energy = 25eV). For shotgun lipidomics the aliphatic constituents were confirmed by product ion scanning (collision energy = 25eV) of fatty acyl carboxylates. Individual molecular species were quantified by comparing the ion intensities of the individual molecular species to that of the lipid class internal standard with additional corrections for type I and type II ^13^C isotope effects. Cardiolipin molecular species were quantified using LC/MS on a Q-Exactive instrument in the negative ion mode. Species and cardiolipin were confirmed by their fatty acyl carboxylates as well as the product ion 153 m/z. Cardiolipins were separated on an Accucore^TM^ C30 column 2.1 x 150 mm (Thermo Scientific) with mobile phase A comprised of 60% acetonitrile, 40% water, 10mM ammonium formate, and 0.1% formic acid and mobile phase B comprised of 90% isopropanol, 10% acetonitrile with 2 mM ammonium formate, and 0.02% formic acid. Initial conditions were 30% B with a discontinuous gradient to 100% B at a flow rate of 0.260 ml/min.

For heatmap visualizations, PL subspecies that were reliably detected more than 1.5% of the total lipid class across all the conditions were retained, followed by visualization using a Volcano plot (GraphPad Prism v10.2.1).

### Gene expression and RNA-seq analysis

Total RNA was extracted from culture cells or frozen tissues using TRIzol (Invitrogen), and reverse transcribed with an iScript cDNA synthesis kit (Bio-Rad). cDNA was quantified using iTaq Universal SYBR Green Supermix (Bio-Rad) on a QuantStudio 6 Flex 384-well qPCR system (Applied Biosystems). Relative mRNA content was evaluated using the delta-delta Ct method, normalized by housekeeping genes (*36B4*). For RNA-seq, RNA was purified using RNeasy Mini Kit (Qiagen) with on-column DNase I digestion (Qiagen). Library preparation and RNA-seq were performed at the UCLA Technology Center for Genomics & Bioinformatics. 200bp paired-end seq cDNA libraries were analyzed with an Agilent TapeStation and sequenced on an Illumina NovaSeq X Plus 10B at 200 cycles.

Raw sequencing reads were demultiplexed using bcl2fastq (Illumina). Fastq files were checked for quality, filtered, and corrected for bp mismatches using fastp (Version 0.23.2) (*80*). Salmon was used with full selective alignment to map the reads to the mouse transcriptome and quantify transcript counts [mm10 - Ensembl – Release 97] (*81*). Raw counts were analyzed using the iDEP2.0 to yield differential gene expressions and PCA analysis (*82*).

For measuring mtDNA abundance, genomic DNA extracted from culture cells or frozen tissues with DNeasy Blood and Tissue Kit (Qiagen) was quantified using iTaq Universal SYBR Green Supermix (Bio-Rad) as described above.

### Electron microscopy

Harvested primary brown adipose tissue were briefly washed with PBS, cut into small pieces (1-2 mm^3^) and immediately fixed with freshly prepared fixative buffer (2% PFA, 2.5% glutaraldehyde, 2.1% sucrose, 100 mM sodium cacodylate) for 72 h at 4°C. Tissues were washed with ice-cold cacodylate buffer (2 mM CaCl_2_, 100 mM sodium cacodylate) 5 times, and incubated with 2% osmium tetroxide, 1.5% potassium ferricyanide in 0.1M sodium cacodylate for 1 hr. The samples were washed 5 times with double-distilled water and incubated in 1% thiocarbohydrazide (TCH) solution for 20 min at RT. Then, these were washed 5 times with double-distilled water and incubated with 2% osmium tetroxide in double-distilled water for 30 min at RT. The samples were washed 5 times with double-distilled water and stained with 2% uranyl acetate overnight at 4°C. After the 5 times wash with double-distilled water, the tissues were dehydrated with increasing concentration of ice-cold ethanol (30%, 50%, 70%, 85%, 95%, 100%, 100%; 8 min each), RT ethanol (100%) 2 times, 8 min each, and infiltrated with 33% Embed812 resin in acetone for 2 h, with 66% Embed812 overnight, and 100% Embed812 for 3 h. The infiltrated tissues were embedded in embedding mold that have been filled to the top with fresh Embed812 resin and polymerized in oven at 60°C for 48 h, and 65 nm sections made with Leica UC6 ultramicrotome were collected onto freshly glow-discharged 100mesh copper grids. The sections were collected and stained with Reynold’s lead citrate. Transmission electron microscopy pictures were taken at California NanoSystems Institute at UCLA.

### Primary brown adipocytes

Primary mouse brown adipocytes (pBA) were prepared as previously described with some modifications (*83*). BAT harvested from 3-4 new-born neonates (P+1) were minced for 30 sec, transferred into 15 ml conical tubes, and incubated with 7.5 mL of digestion buffer (100 mM HEPES pH 7.4, 123 mM NaCl, 1.3 mM CaCl_2_, 5 mM KCl, 5 mM glucose, 2% essentially fatty acid-free BSA, 2mg/ml type 2 collagenase (Worthingon)) for 40 min at 37°C under inverted rotation. The digested BAT suspension was then adjusted to 15 mL with ice-cold Dulbecco’s modified Eagle’s medium (DMEM: Corning 10-013-CM) and gently passed through an 18.5G needle (8 times) for extra disruption. Digested tissue was strained with 40 µm filters into 50 mL conical tubes, followed by washing the filters with 30 mL ice-cold DMEM and spun down at 600 × *g* for 10 min. The collected stromal vascular cell (SVC) fraction were resuspended with 30mL ice-cold DMEM, spun down at 600 × *g* for 10 min, resuspended with warm pBA growth medium (DMEM supplemented with 15% Fetal Bovine Serum (GemCell^TM^), Penicillin (50 U/ml), and streptomycin (50 µg/ml)) and plated in 10 cm dishes. After 48 hours, cells were rinsed and cultured with fresh pBA growth medium. Once cells reached ∼80% confluence, they were subcultured to experiment format and left until they become over-confluent. For the differentiation, SVC were grown to confluence and treated with rosiglitazone (1 µM), indomethacin (125 µM), 3-isobutyl-1-methylxanthine (0.5 mM), dexamethasone (1 µM), Humulin (100nM) and triiodothyronine (2 nM) for 48 h (day 0-2), rosiglitazone (1 µM), Humulin (100nM) and triiodothyronine (2 nM) for 48 h (day 2-4), and Humulin (100nM) and triiodothyronine (2 nM) for 48 h (day 4-6). For RNA isolation, the differentiation medium was replaced with fresh pBA growth medium and stimulated with 1 µM norepinephrine (Baxter) for 4 h.

### Live cell imaging

Primary mouse brown adipocytes were differentiated in 35 mm glass-bottom dishes (P35GC-1.5-14-C, MatTek). On the day 6 of differentiation, cells were incubated with 100 nM Mitotracker Green (Thermo Fisher Scientific) and 100 nM LipidTOX Red (1:1000) (Thermo Fisher Scientific) in pBA growth medium at 37°C for 15 min, washed-out and destained by incubation with fresh pBA growth medium at 37°C for 45 min.

### Seahorse respirometry

At day 4 of differentiation, primary brown adipocytes were subcultured with 1 mg/ml type 2 collagenase (Worthingon) in STEMPro accutase (ThermoFisher) and seeded on 24-well Seahorse cell culture plate with 15 x 10^3^ cells/well in 50 µl of differentiation medium (rosiglitazone (1 µM), Humulin (100 nM) and triiodothyronine (2 nM)). The next day, additional 250 µl of differentiation medium (rosiglitazone (1 µM), Humulin (100 nM) and triiodothyronine (2 nM)) was added to each well. On day 6 of differentiation, cells were washed with Seahorse incubation medium (1 x DMEM (D5030, Sigma), 5 mM HEPES (pH 7.4), 32 mM NaCl, 0.6% phenol red (P0290, Sigma), 1% essentially fatty acid-free BSA, pH adjusted to pH 7.4) twice and 450 µl Seahorse incubation medium was added to the well. Oxygen consumption rates (OCR) were measured using a Seahorse XFe24 Analyzer (Agilent). During the run, cells were sequentially treated with norepinephrine (1 µM), oligomycin (4 µM), FCCP/pyruvate (10 µM/ 500 µM), and rotenone/antimycin A (7.5 µM/4 µM). After the measurements, cells were washed with PBS (+Ca^2+^ and Mg^2+^) 4 times and lysed with NaOH lysis buffer (0.33 M NaOH, 0.1% SDS) for the protein determination.

For evaluating respiratory capacity through individual ETC complex, primary brown adipocytes were permeabilized by 3 nM Seahorse XF Plasma Membrane Permeabilizer (Agilent) in KCl-MAS (100 mM KCl, 10 mM KH_2_PO_4_, 2 mM MgCl_2_, 1 mM EGTA, 5 mM HEPES (pH7.4) with 0.1% essentially fatty acid-free BSA, pH adjusted to pH 7.4)) supplemented with individual substrates/inhibitors. For beta oxidation activity, which requires CI, CII, ETF, CIII and CIV, cells were incubated with malate (0.5 mM) followed by sequentially treated with palmitoyl carnitine/ADP (40 µM/4 mM) and rotenone/antimycin A (2 µM/4 µM). For CII-CIII-CIV activity, permeabilized cells were sequentially treated with succinate/ADP/rotenone (5 mM/4 mM/2 µM) and antimycin A (4 µM). For CIV activity, permeabilized cells were sequentially treated with TMPD/ascorbate (500 µM /700 µM) and sodium azide (33 mM).

Respiratory capacity in isolated BAT mitochondria was evaluated on Seahorse XF96 Analyzers (Agilent). Two lobes of BAT from one mouse were transferred to 2 ml glass Dounce homogenizer containing 1.2 ml isolation buffer (200 mM sucrose, 10 mM Tris-HCl pH 7.4, 1 mM EGTA, 20 mM HEPESs) supplemented with 0.2% essentially fatty acid-free BSA, and homogenized with 20 strokes using a loose pestle (pestle A) and 20 strokes using a tight pestle (pestle B). The resulting homogenates were transferred into 2 ml Eppendorf tubes. The glass homogenizers were rinsed with an additional 0.8 ml isolation buffer and the rinse was combined with homogenates in the 2 ml Eppendorf tubes. The homogenates were centrifuged at 600 × *g* for 10 min at 4°C. Subsequently, 1.5 ml of the supernatants were transferred to new 1.5 ml Eppendorf tubes and centrifuged at 7,000 × *g* for 10 min at 4°C. The pellets were washed with 1.5 ml isolation buffer and centrifuged again at 7,000 × *g* for 10 min at 4°C. The final pellets were resuspended in 50 µl of isolation buffer, and protein concentrations were determined using the BCA Protein Assay Kit (Pierce). The protein concentration was adjusted to 10-20 mg/ml, and equal amounts of mitochondria derived from 4-5 mice were pooled. Isolated BAT mitochondria were loaded in an XF96 microplate at 3.5 µg per well for CI, CII, and long-chain fatty acid oxidation measurements and 1 µg per well for CIV. Mitochondria were plated in KCL buffer (115 mM KCl, 10 mM KH_2_PO_4_, 2 mM MgCl_2_, 5 mM HEPES, 1 mM EGTA, 0.1% fatty acid-free BSA, pH adjusted to pH 7.2) with 1 mM GDP and centrifuged at 2100 xg for 10 minutes at 4°C. With all substrates, state 3 respiration was measured with 4 mM ADP. Long-chain fatty acid oxidation was measured with 40 µM palmitoylcarnitine and 0.5 mM malate, Complex I with 5 mM pyruvate and 1 mM malate, and Complex II with 5 mM succinate and 2 µM rotenone. Injections included: 4 µM oligomycin, 4 µM FCCP, and rotenone/antimycin A (2 µM/4 µM). For Complex IV activity, mitochondria were plated with 4 µM antimycin A followed by injection with 0.5 mM TMDP/1 mM ascorbate, and then 50 mM azide to inhibit CIV.

### Live-cell labeling with fatty acid-based clickable photoaffinity probes

On day 3 of differentiation, primary brown adipocytes (pBA) were subcultured with 1 mg/ml type 2 collagenase (Worthingon) in STEMPro accutase (ThermoFisher) and seeded in 6-well plate at the density of 1.5 x 10^6^ cells/well. On day 6 of differentiation, cells were washed once with DMEM and treated with the indicated photoaffinity probe (20 µM) in DMEM supplemented with 2% essentially fatty acid-free BSA at 37°C for 1 h. For HTS-3 treatment, cells were pretreated with HTS-3 (10 µM) in pBA growth medium for 2 h and exposed to a photoaffinity probe with HTS-3 (10 µM) in DMEM supplemented with 2% essentially fatty acid-free BSA. After incubation, cells were washed twice with PBS and were directly irradiated with 365 nm light at 100 mJ/cm^2^ for 10 min using UVP Crosslinker CL-1000 (AnalytikJena). After UV cross linking, cells were harvested with cold DPBS and washed with cold DPBS twice by centrifugation (3000rpm, 3min). The isolated cell pellets were lysed by probe sonication in 300 µL DPBS and were centrifuged (100,000 × *g*, 45 min, 4°C). The resulting membrane fraction (pellet) was resuspended in 200 µL DPBS by probe sonication and was stored at –80°C.

### Gel-based visualization of probe-labelled proteins

The prepared crosslinked membrane fraction was diluted to 1.0 mg/ml, and 50 µL of the proteome was transferred to new Eppendorf 1.5 mL tube. The proteome was incubated with 6 µL of click reagent containing CuSO4 (1.0 µl/sample, 50mM in H_2_O), TBTA (3.0 µl/sample, 1.7 mM in 4:1 DMSO:*t*-BuOH), Alexa 488-N_3_ (1.0 µl/sample, 1.25mM in DMSO), and TCEP (1.0 µl/sample, 50mM in H_2_O). The reaction was carried out for 1 h at room temperature with vortexing every 15 min and quenched by adding 4 x SDS sample buffer (15 µl). The samples were resolved by SDS-PAGE (NuPAGE 4-12% Bis-Tris gels (Invitrogen)) and visualized on a Bio-Rad ChemiDoc with Alexa Fluor 488 filter.

### Streptavidin enrichment of probe-labelled proteins

The prepared crosslinked membrane fraction was diluted to 1.0 mg/ml, and 250 µL of the proteome was transferred to new Eppendorf 1.5 mL tubes. The proteome was incubated with 30 µL of click reagent containing CuSO4 (5 µl/sample, 50 mM in H_2_O), TBTA (15 µl/sample, 1.7 mM in 4:1 DMSO:*t*-BuOH), *N*-(3-Azidopropyl)biotinamide (5 µl/sample, 10 mM in DMSO, A2524 (TCI)), and TCEP (5 µl/sample, 50 mM in H_2_O) for 1 h at room temperature with vortexing every 15 min. Following the CLICK reaction, 10% SDS (25 µl/sample), SP3 beads (50 µl/sample, 1:1 GE45152105050250:GE65152105050250 (Cytiva)) were added and incubated at RT for 10 min with vortexing at 1000 rpm. Proteins were bound to SP3 beads by adding 100% ethanol (400 µl/sample) followed by vortexing at 1000 rpm at RT for 10 min. The SP3 beads were washed with 80% ethanol (400 µl x 3) and proteins were eluted with 0.2% SDS in PBS (50 µl x 2). The eluents (100 µl/sample) were subsequently incubated with 500 µl of 5% streptavidin-agarose beads (Pierce) for 2 h at RT. The beads were washed sequentially with 2 x 1 ml 0.1% SDS in PBS, 1 ml 1 M KCl, 1 mL 0.1 M Na_2_CO_2_, 1 mL 2 M urea in PBS, 2 x 1 mL PBS. After washing, the beads were pelleted by centrifugation (1,500 × *g*, 2 min) and PBS was removed. The proteins were eluted from beads by boiling with 4 x sample buffer containing 2 mM biotin (50 µl/sample) for western blot analysis.

### Proteomics analysis of probe-labelled proteins

Immunoprecipitated streptavidin beads were resuspended in 100 μl lysis buffer (12 mM sodium lauroyl sarcosine, 0.5% sodium deoxycholate, 50 mM triethylammonium bicarbonate (TEAB), Halt™ Protease, and Phosphatase Inhibitor Cocktail) and subjected to bath sonication (10 min, Bioruptor Pico, Diagenode Inc.; Denville, NJ). Samples were treated with 10 μl tris (2-carboxyethyl) phosphine (55 mM in 50 mM TEAB, 30 min at 37 °C), followed by treatment with 10 μL chloroacetamide (120 mM in 50 mM TEAB, 30 min at 25 °C in the dark). Samples were then diluted 5-fold with aqueous 50 mM TEAB and incubated overnight with Sequencing Grade Modified Trypsin (1 μg in 10 μl 50 mM TEAB; Promega, Cat # V511A, Madison, WI, USA). Following digestion, equal volumes of ethyl acetate/trifluoroacetic acid (TFA, 100/1, *v/v*) were added, after which samples were mixed (5 min), and subjected to centrifugation (13,000 × *g*, 5 min). The upper phase was discarded and the lower phase was dried in a centrifugal vacuum concentrator. Dried samples were reconstituted in acetonitrile/water/TFA (solvent A, 100 μL, 2/98/0.1, *v/v/v*) and loaded onto a small portion of a C18-silica disk (3M, Maplewood, MN, USA) placed in a 200 μl pipette tip. Prior to sample loading, the C18 disk was prepared by the sequential treatment with methanol (20 μl), acetonitrile/water/TFA (solvent B, 20 μl, 80/20/0.1, *v/v/v*), and solvent A (20 μl). After loading the sample, the disc was washed with solvent A (20 μl, eluent discarded) and eluted with solvent B (40 μl). The collected eluent was dried in a centrifugal vacuum concentrator.

The eluents were dried and reconstituted in water/acetonitrile/FA (solvent B, 10 μl, 98/2/0.1, *v/v/v*), and the aliquots (5 μl) were injected onto a reverse-phase nanobore HPLC column (AcuTech Scientific, C18, 1.8 μm particle size, 360 μm × 20 cm, 150 μm ID, San Diego, CA, USA), equilibrated in solvent E, and eluted (500 nl/min) with an increasing concentration of solvent F (acetonitrile/water/FA, 98/2/0.1, *v/v/v*: min/% F; 0/0, 5/3, 18/7, 74/12, 144/24, 153/27, 162/40, 164/80, 174/80, 176/0, 180/0) using an Eksigent NanoLC-2D system (Sciex, Framingham, MA, USA). The effluent from the column was directed to a nanospray ionization source connected to a hybrid quadrupole-Orbitrap MS (Q Exactive Plus, ThermoFisher Scientific, Waltham, MA, USA), acquiring mass spectra in a data-dependent mode alternating between a full scan (350–1700 m/z, automated gain control (AGC)) target 3×10^6^, 50 ms maximum injection time, FWHM resolution 70,000 at 200 m/z) and up to 15 MS/MS scans (quadrupole isolation of charge states 2–7, isolation window 0.7 m/z) with previously optimized fragmentation conditions (normalized collision energy of 32, dynamic exclusion of 30 s, AGC target 1×10^5^, 100 ms maximum injection time, FWHM resolution 35,000 at 200 m/z).

The raw proteomic data were queried against the UNIPROT mouse-reviewed protein database using SEQUEST-HT in Proteome Discoverer (Version 2,4, Thermo Scientific, Waltham, MA, USA), which provided measurements of abundances for the identified peptides in each sample normalized to total protein amount. Decoy database searching was used to identify high confidence tryptic peptides (FDR < 1%). Tryptic peptides containing amino acid sequences unique to individual proteins were used to identify and provide the relative quantification between proteins in each sample. The median abundance values of all replicates from each condition were used to generate abundance ratios for each protein across the groups.

### Blue Native Page

#### Mitochondria isolation

Tissues or cells were transferred to 2 ml glass Dounce homogenizer containing 1.2 ml isolation buffer (200 mM sucrose, 10 mM Tris-HCl pH 7.4, 1 mM EGTA, 20 mM HEPES, protease inhibitors) supplemented with 0.2% essentially fatty acid-free BSA, and homogenized with 20 strokes using a loose pestle (pestle A) and 20 strokes using a tight pestle (pestle B). The resulting homogenates were transferred into a 2 ml Eppendorf tubes. The glass homogenizers were rinsed with an additional 0.8 ml isolation buffer and the rinse was combined with homogenates in the 2 ml Eppendorf tubes. The homogenates were centrifuged at 600 × *g* for 10 min at 4°C. Subsequently, 1.5 ml of the supernatants were transferred to new 1.5 ml Eppendorf tubes and centrifuged at 7,000 × *g* for 10 min at 4°C. The pellets were washed with 1.5 ml isolation buffer and centrifuged again at 7,000 × *g* for 10 min at 4°C. The final pellets were resuspended in 150 µl of isolation buffer containing 10% glycerol, and protein concentrations were determined using the BCA Protein Assay Kit (Pierce). The isolated mitochondria were aliquoted at 100 µg protein/tube and stored at –80°C until further analysis.

#### Running blue native page

Blue native page was performed as previously described with some modifications (*84*). Fifty microgram of mitochondria suspension was spun down (7,000 × *g*, 10 min, 4°C) to remove glycerol-containing buffer. The resulting pellet was resuspended in 20 µl of 1x Sample Buffer (Invitrogen) with 2% digitonin and was incubated on ice for 20 min. After centrifugation (20,000 × *g*, 10 min, 4°C), 15 µl of the supernatant was mixed with 2 µl of Coomasie G-250 (5%). Samples were loaded onto NativePAGE 3-12% Bis-Tris gels (Invitrogen) and electrophoresed at 4°C in constant voltage at 150 V for 36 min with Dark Blue Cathode Buffer (contains 0.02% G-250), followed by at 250 V for 150 min with Light Blue Cathode Buffer (contains 0.002% G-250). For the membrane blotting, electrophoresed proteins were transferred onto Amersham Hybond P 0.45 µm PVDF membranes (Cytiva) (25 mV for 60 min). The proteins were fixed on the membrane with 8% acetic acid for 15 min, air-dried, destained by methanol wash for 5 min, and blotted as described before. For in-gel activity assay, gels were incubated with Complex IV substrate solution (5 mg Diaminobenzidine, 10 mg cytochrome c, 4.5 ml phosphate buffer pH 7.4 (100mM), 750 mg sucrose to 5.5 ml H_2_O) or Complex I substrate solution (2 mg NADH, 50 mg Nitrotetrazolium Blue chloride, 40 µl Tris-HCl pH 7.4 (1 M) to 20 ml H_2_O). The enzymatic reactions were stopped by adding 10% acetic acid, and stored in water at 4°C.

### Vector Construction

Full-length LPCAT3 (NCBI RefSeq: NM_145130.2) was PCR amplified from *Mus musculus* C57BL/6J interscapular BAT cDNA. The N-terminal FLAG epitope-tag and Gateway overhangs were added via primer design. Flag-Lpcat3 was cloned into Gateway pDONR^TM^221 entry vector (BP Clonase^TM^ II, ThermoFisher). The catalytically inactive (His-374-Ala) mutant LPCAT3 was generated by site-directed mutagenesis. All sequences were subcloned into the pLEX_307 (SV40-Puro; EF1a-gateway-V5 tag) lentiviral destination vector (a kind gift from David Root, Addgene plasmid # 41392) by attL x attR recombination (LR Clonase^TM^ II Enzyme mix, ThermoFisher).

### Lentivirus Generation

Lentivirus was generated in 293T cells by transfecting lentiviral vectors with pMDL/pRRE, pRSV-Rev, and pMD2.VSVG packaging vectors. The supernatant was collected 48 and 72 h after transfection, passed through a 0.45 µm filter (Millipore-Sigma), and collected into an open-top thin wall ultra-clear tube (Beckman, #344058). Ice-cold 15% sucrose (Millipore-Sigma) was added on top of the supernatant and centrifuged (Beckman Coulter Optima XPN-100) at 22,000 rpm at 4°C for 2.5 h. The lentiviral pellets were gently re-dissolved in PBS and frozen at –80°C.

For lentiviral transduction in primary brown adipocytes, undifferentiated cells were seeded at a density of 5 x 10^4^ cells/ml. The next day, cells were infected with lentivirus carrying mCherry, LPCAT3^WT^, or LPCAT3^H374A^ in the presence of 8 μg/ml polybrene (Millipore-Sigma). Sixteen hours later, medium was replaced with fresh culture media. Five days post-transduction, differentiation was started. Despite the avoidance of drug selection due to its adverse impact on differentiation, over 80% mCherry transgenes were confirmed microscopically.

### Patch-clamp recordings

Patch-clamp recording was performed from isolated brown fat mitoplasts of mice as previously described(*3*). The mitoplasts used for patch-clamp experiments were 3 to 5 μm in diameter and typically had membrane capacitances of 0.3 to 1.2 pF. Both the bath and pipette solutions were formulated to record H^+^ currents and contained only salts that dissociate into large anions and cations that are normally impermeant through ion channels or transporters (pH 7.0).

Pipettes were filled with 130 mM tetramethylammonium hydroxide, 1 mM EGTA, 2 mM Tris–HCl, and 100 mM Hepes. pH was adjusted to 7.0 with d-gluconic acid, and tonicity was adjusted to ∼360 mmol/kg with sucrose. Typically, pipettes had resistances of 25 to 35 MΩ, and the access resistance was 40 to 75 MΩ. Whole-mitoplast H^+^ current was recorded in the bath solution containing 100 mM Hepes and 1 mM EGTA (pH adjusted to 7.0 with Trizma base, and tonicity adjusted to ∼300 mmol/kg with sucrose). All experiments were performed under continuous perfusion of the bath solution. All electrophysiological data presented were acquired at 10 kHz and filtered at 1 kHz.

### Coarse grain molecular dynamics simulations (CG-MD)

The initial structures for the *Mus musculus* complexIV and VDAC1 were obtained from the Protein Data Bank (https://www.rcsb.org/) with PDB ID 7o3e (*48*) and PDB ID 4c69 (*85*), respectively. For the Ant2 protein (UniProt ID P51881) and MTCH2 (UniProt ID Q9Y6C9), the corresponding sequences for *M. musculus* deposited in UniProt (https://www.uniprot.org/) were used for structural modeling with AlphaFold2 (AF2) (*86*).

The three systems were subjected to a vacuum all-atom minimization stage for a maximum of 50,000 steps or until the maximum force on any atom was less than 100 kJ mol^−1^ nm^−1^. The AMBER99SB-ILDN force field (*87*) and the steepest descent algorithm were employed for this procedure. Once the initial structure of each protein was minimized, they were converted to CG model using *Martinize2* (https://doi.org/10.7554/eLife.90627.1). All the CG proteins were embedded in a DOPC:Cardiolipin:SAPE:DOPE membrane with molar a ratio of 50:25:12.5:12.5, respectively, using the *INSert membrANE* program (*88*). The systems were solvated with a ∼0.15 M NaCl solution.

The CG-MD simulations were carried out using the GROMACS 2023.3 software (https://doi.org/10.1016/j.softx.2015.06.001), and the Martini 3 force field (*89*). To maintain the 3D structure of the proteins, an elastic network was applied with a force constant of 500 kJ mol^−1^ nm^−2^. Temperature was controlled by the V-rescale thermostat at 310 K, with a coupling constant of 1 ps (*90*). For the pressure, the Parrinello-Raman barostat (https://doi.org/10.1063/1.328693) was employed with a reference pressure of 1 bar and a coupling constant of 12 ps. Two replicates with different initial velocities for 10 μs were carried out for each system. A time step of 20 fs was used.

For the 2D lipid preference analysis, density maps were obtained using the gmx_densmap tool with a bin size of 0.05 was set. The difference between the PUFA and MUFA densities (integrated along the z-axis) at each point on the (x,y) map was plotted.

For the analysis of protein-PUFA interaction, the contact frequency between PUFA and the proteins was obtained using the gmx_select tool; A cutoff of 0.5 nm was selected to consider a protein-lipid contact.

For all analysis, the first 2 μs were omitted to guarantee the correct mixing of lipids around the proteins.

### Evaluation of Mitochondrial Membrane Fluidity

Isolated mitochondria were diluted to a final concentration of 0.8 µg/µL in isolation buffer (200 mM sucrose, 10 mM Tris-HCl pH 7.4, 1 mM EGTA, 20 mM HEPES, 10% glycerol). The diluted mitochondria were then stained with 10 µM C-Laurdan and incubated for 5 minutes. General polarizability was assessed over a range of temperatures using a Cary Eclipse Fluorescence Spectrometer equipped with a Cary Temperature Controller.

Samples were excited at 352 nm with a slit width of 10 nm, and the emission was measured at 440 nm and 490 nm using a slit width of 2.5 nm. The samples were initially cooled to 3°C then the temperature was then gradually increased at a rate of 0.5°C/min, and spectral intensity was measured in steps at each 1°C increment until the temperature reached 42°C. General polarizability was calculated by the following formula: 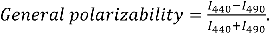

### Statistical Analysis

All data are presented as mean ± SEM. The number of mice in each study is reported in the figure legends.

Statistical analyses were performed in Prism 10 software (GraphPad) using two-tailed Welch’s *t*-test for parametric comparisons between two groups and two-way ANOVA for comparisons of multiple groups. If a parameter was measured over time, data were analyzed by repeat measures two-way ANOVA with time as an independent factor. *P* values < 0.05 were considered significant and presented as *p*<0.05, *p*<0.01, or *p*<0.001. No statistical method was used to predetermine sample size.

## Supplementary Figure

**Fig. S1.**
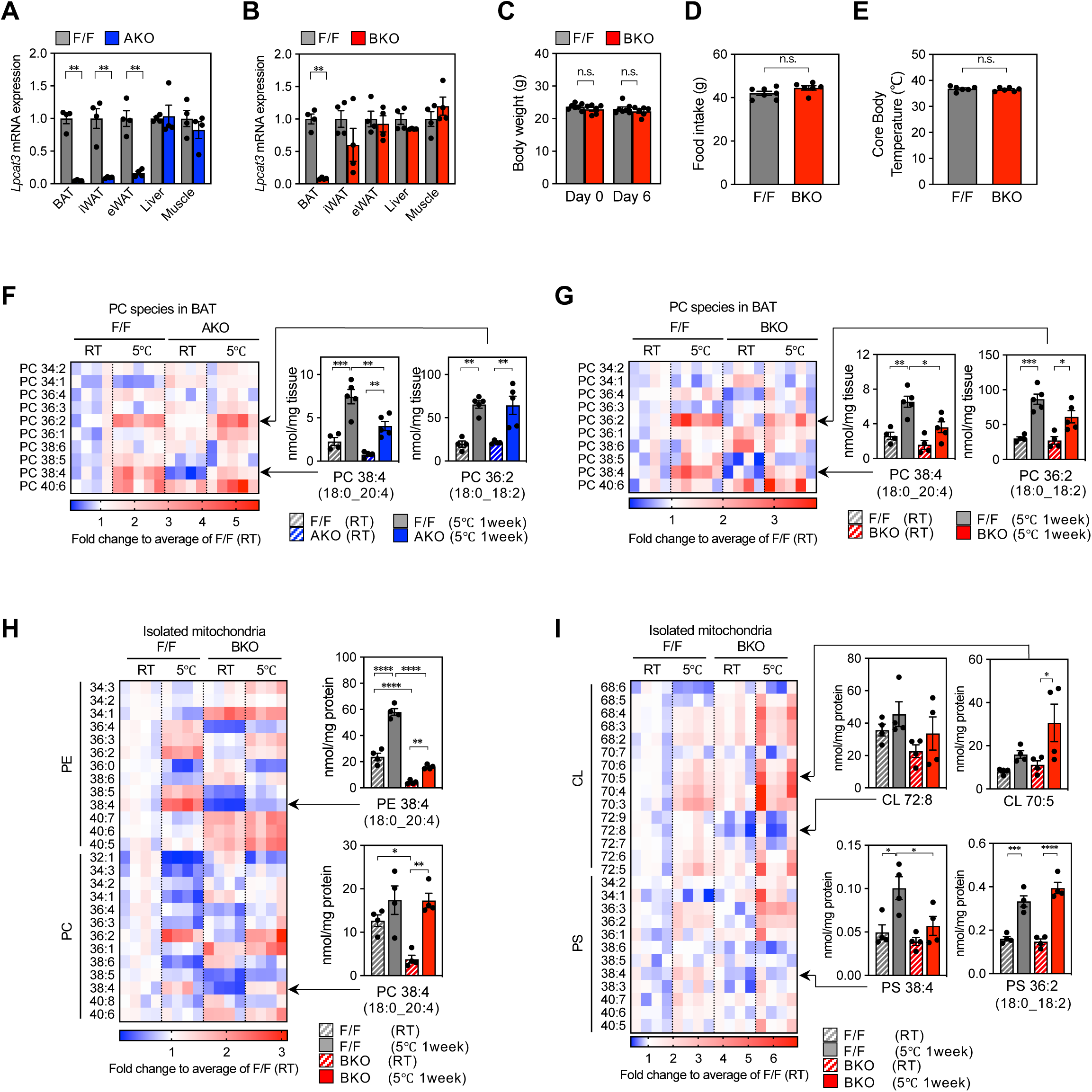
Cold-inducible AA-PE in brown fat mitochondria requires LPCAT3 activity. (**A, B**) qPCR analysis of *Lpcat3* transcript levels in fat depots, liver, and quadriceps muscle from (**A**) F/F and *Lpcat3*^AKO^ mice, or (**B**) F/F and *Lpcat3*^BKO^ mice housed at 23℃ (*n* = 4/group). (**C-E**) Consequence of cold adaptation on (**C**) body weight, (**D**) food intake, and (**E**) core body temperature of F/F and *Lpcat3*^BKO^ mice (*n* = 6, 7). Mice were transferred from 23℃ to 5℃ for 7 days, with *ad libitum* access to food and water. (**F, G**) LC-MS/MS analysis of PC composition in BAT from (**F**) F/F and *Lpcat3*^AKO^ mice, or (**G**) F/F and *Lpcat3*^BKO^ mice housed at either 23℃ or 5℃ for 7 days (*n* = 4–5/group). 38:4-PC and 36:2-PC subspecies are depicted in individual bar graphs. (**H, I**) LC-MS/MS analysis of (**H**) PC/PE and (**I**) PS/CL molecular species in isolated BAT mitochondria from F/F and *Lpcat3*^BKO^ mice housed at either 23℃ or 5℃ for 7 days (*n* = 4, 4). Data are presented as mean ± SEM. \**p*<0.05, \*\**p*<0.01, \*\*\**p*<0.001, \*\*\*\**p*<0.0001 by two-sided Welch’s *t*-test (A, B, C, D, E), or one-way ANOVA with Tukey’s multiple comparisons test (F, G, H, I).

**Fig. S2.**
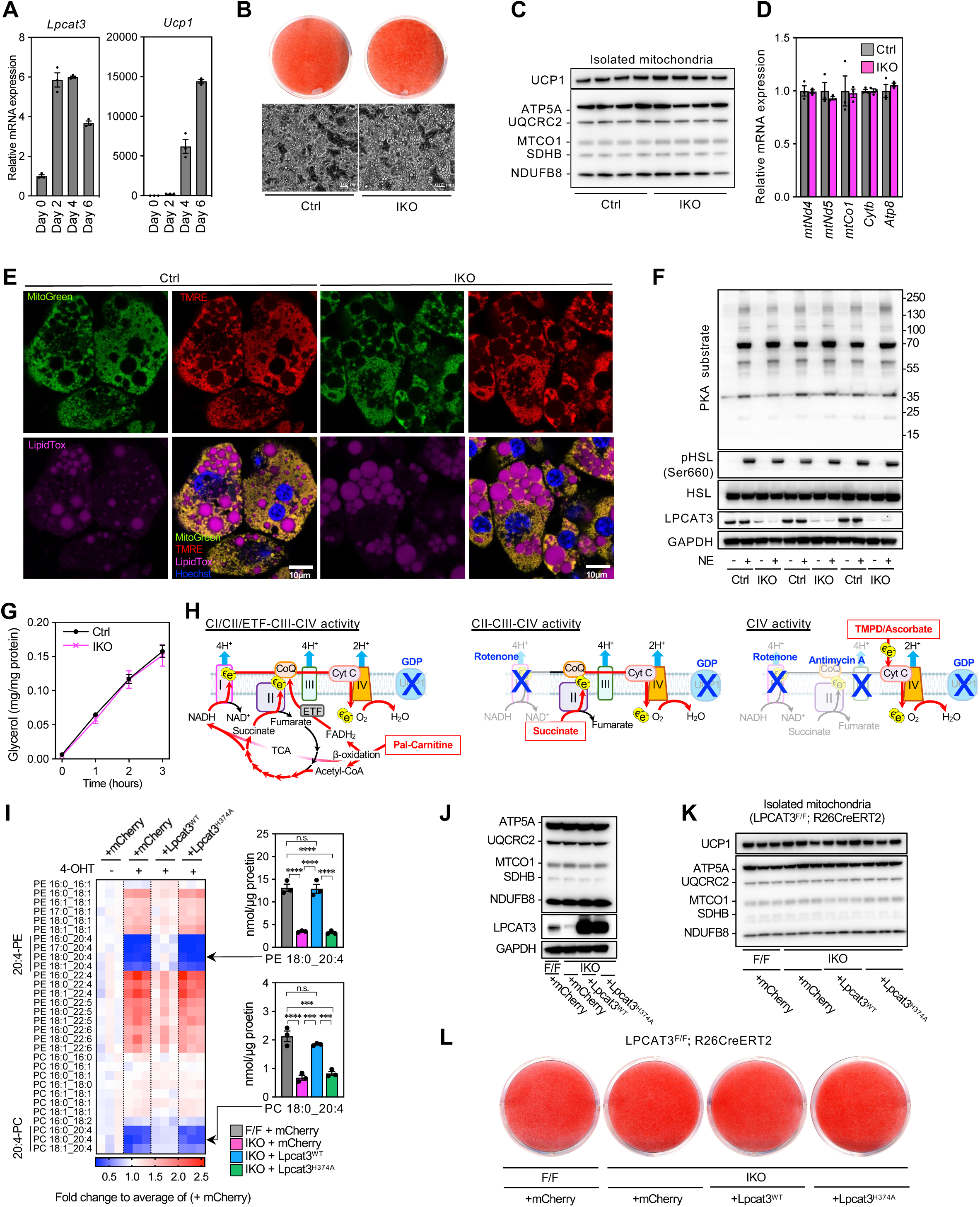
Analysis of 4-OHT-inducible *Lpcat3*^IKO^ pBAs. (**A**) qPCR analysis of *Lpcat3* and *Ucp1* transcript levels in pBAs over the course of differentiation (*n* = 3). (**B**) Oil red-O staining and bright-field microscopy analysis of *Lpcat3*^F/F^, *Cre*^ERT2^ pBAs treated with either vehicle (Ctrl) or 4-OHT (*Lpcat3*^IKO^) and differentiated for 6 days. (**C**) Western blot analysis of UCP1 and OXPHOS subunits in isolated mitochondria from Ctrl and *Lpcat3*^IKO^ pBAs (day 6 of differentiation, *n* = 4). (**D**) qPCR analysis of mtDNA-encoded gene transcripts in Ctrl and *Lpcat3*^IKO^ pBAs (day 6 of differentiation, *n* = 3). (**E**) Live-cell imaging of Ctrl and *Lpcat3*^IKO^ pBAs stained with MitoTracker-Green, TMRE, and LipidTOX (scale bar, 10 μm). (**F**) Western blot analysis of lipolytic markers in Ctrl or *Lpcat3*^IKO^ pBAs treated with or without NE (1 μM) for 15 min (*n* = 3). (**G**) Lipolytic activity in Ctrl and *Lpcat3*^IKO^ pBAs (*n* = 4). Glycerol release was monitored over time (0–3 h) following treatment with NE (1 μM). (**H**) Graphic illustration of measuring OXPHOS complex-specific activities. (**I**) Lipidomic analysis of PE/PC species in Ctrl pBAs stably expressing mCherry and *Lpcat3*^IKO^ pBAs reconstituted with mCherry, FLAG-LPCAT3^WT^ or FLAG-LPCAT3^H374A^ (day 6 of differentiation, *n* = 3/group). 18:0_20:4-PE and 18:0_20:4-PC are depicted in individual bar graphs. (**J-L**) Western blot analysis of (**J**) LPCAT3 protein levels and OXPHOS subunits in whole-cell lysates, (**K**) UCP1 protein levels and OXPHOS subunits in purified mitochondria, and (**L**) Oil-red-O staining of Ctrl pBAs expressing mCherry and *Lpcat3*^IKO^ pBAs stably reconstituted with mCherry, FLAG-LPCAT3^WT^ or FLAG-LPCAT3^H374A^. GAPDH was used as a loading control. Data are presented as mean ± SEM. \*\*\**p*<0.001, \*\*\*\**p*<0.0001 by one-way RM ANOVA with Tukey’s multiple comparisons test (I).

**Fig. S3.**
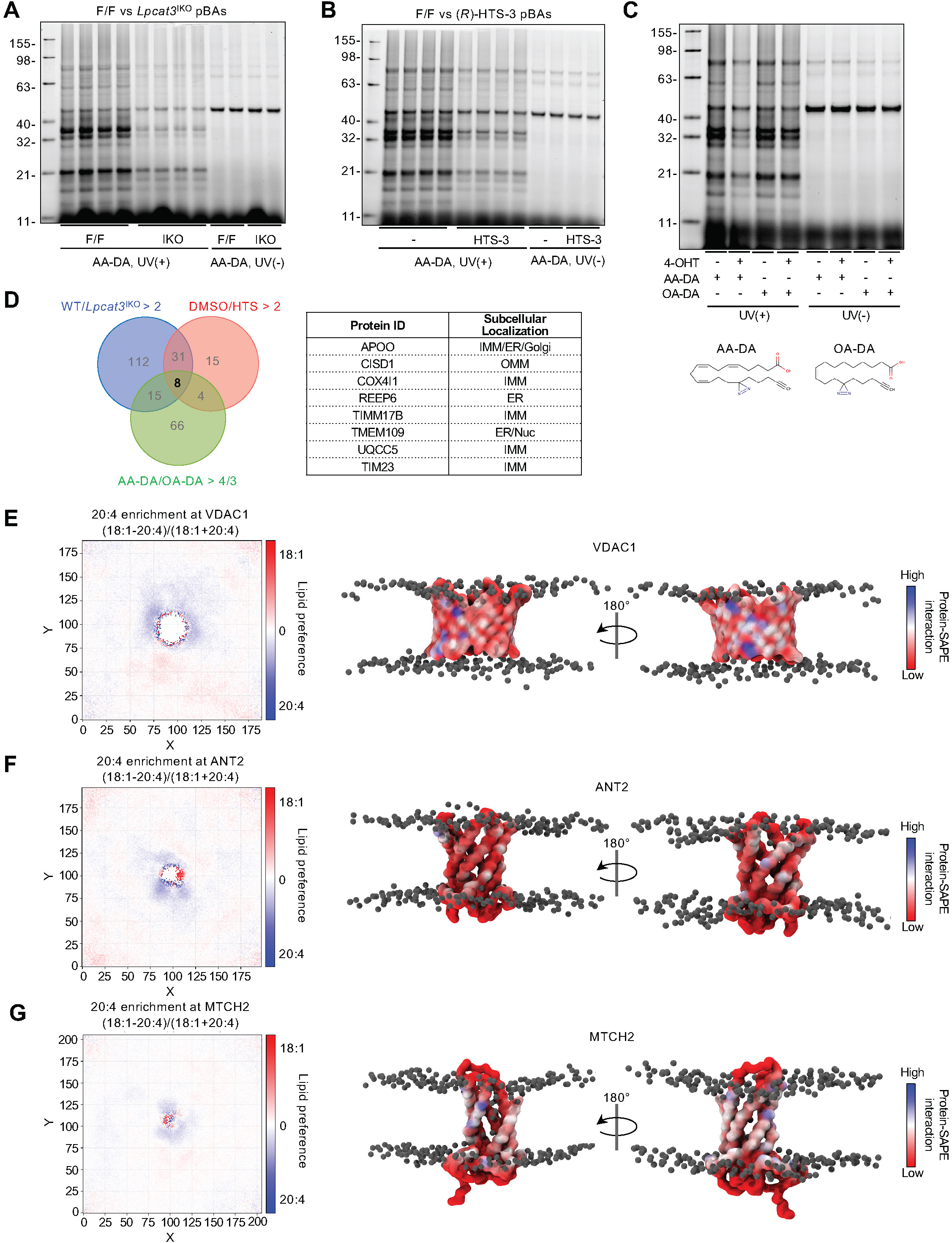
Analysis of arachidonoyl phospholipid-interacting proteins in brown adipocytes. (**A-C**) Evaluation of the phospholipid–protein interactome in pBAs. (**A**) Ctrl and *Lpcat3*^IKO^ pBAs, or (**B**) Ctrl pBAs pre-treated with DMSO or the LPCAT3 inhibitor (*R*)-HTS-3 (10 µM) were incubated with AA-DA (20 µM), UV-irradiated, and the crosslinked proteome was detected by *in*–*gel* fluorescence scanning. (**C**) Ctrl and *Lpcat3*^IKO^ pBAs were differentiated for 6 days, followed by incubation with AA-DA (20 µM) or OA-DA (20 µM), UV-irradiated, and crosslinked proteome was detected by *in–gel* fluorescence scanning. Non-UV-irradiated (UV(-)) groups served as a negative control. (**D**) Left: Venn diagram illustrating the overlap between AA-DA-labeled proteins enriched in (*1*) Ctrl *vs*. *Lpcat3*^IKO^ pBAs, (*2*) DMSO *vs*. (*R*)-HTS-3 treated pBAs, and (*3*) AA-DA vs. OA-DA treated pBAs, identified by streptavidin pull-down and in a UV-dependent manner (>5 fold relative to UV(-)). Right: A table listing ID and subcellular localization of 8 proteins that satisfy all three criteria. (**E-G**) Right: 2D lipid preference analysis by coarse-grain molecular dynamics (CG–MD) simulations on (**E**) VDAC1, (**F**) ANT2 complexes, and (**G**) MTCH2 embedded in a virtual bilayer (comprised of 18:0/20:4-PE, 12.5%; 18:1/18:1-PE, 12.5%; 18:1/18:1-PC, 50%; and tetra-18:2-CL, 25%). The area highly enriched in 20:4-over 18:1-containing phospholipids is shown in blue. Left: analysis of protein-PUFA interaction; contact frequency between 20:4 and the proteins. The images show side views of VDAC1 (E), ANT2 (F), or MTCH2 (G) embedded in the membrane, with the area interacting with PUFA chains shown in blue.

**Fig. S4.**
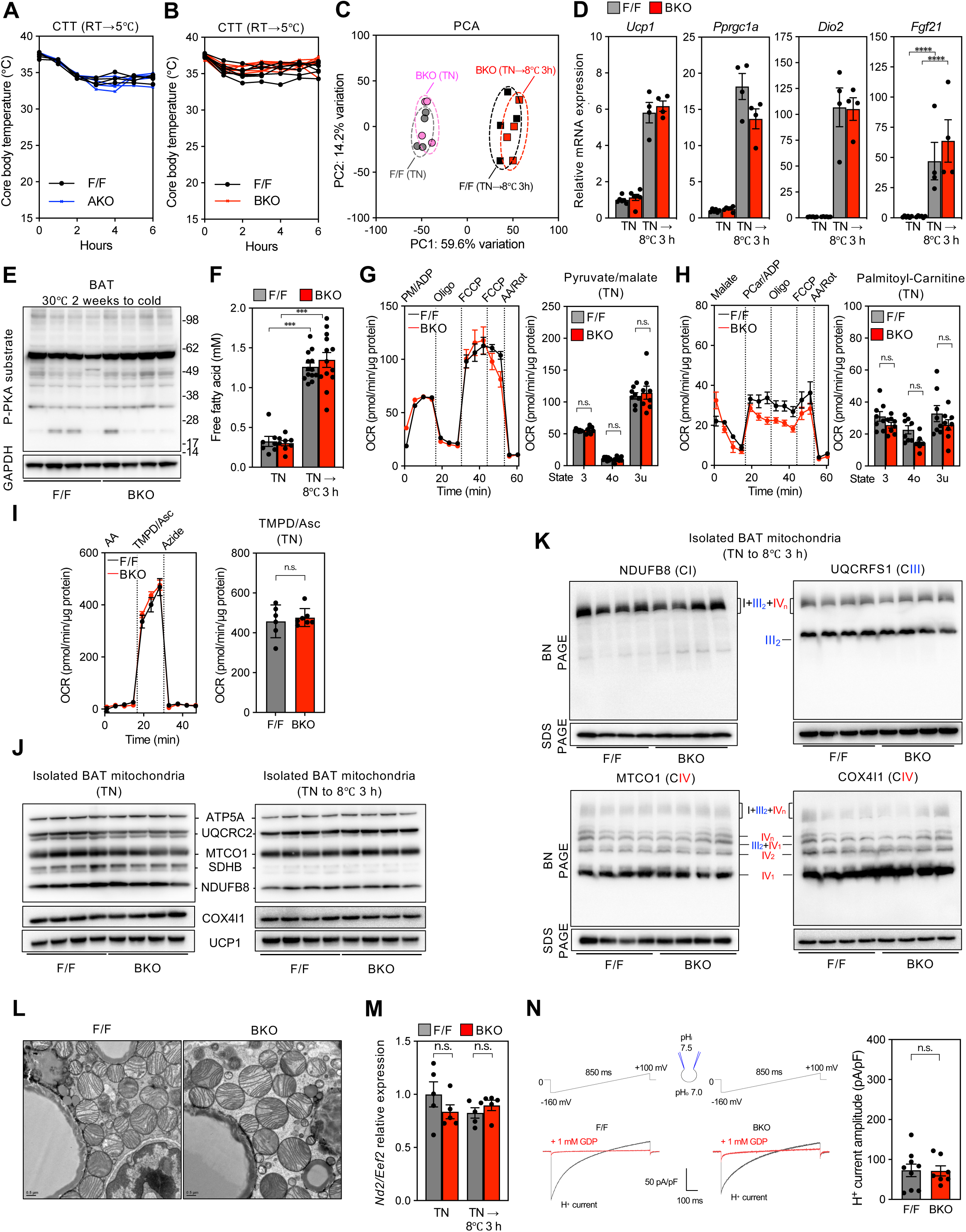
LPCAT3 deficiency in BAT does not affect adrenergic signaling and UCP1 function. (**A, B**) Cold tolerance tests (CTT) were performed 10-week-old male (**A**) F/F and *Lpcat3*^AKO^ mice, or (**B**) F/F and *Lpcat3*^BKO^ mice (*n* = 7/group). Mice were transferred directly from 23℃ to 5℃, followed by hourly monitoring of core body temperature. (**C**) Principal component analysis (PCA) of the BAT transcriptome. RNA-seq was performed on BAT from F/F and *Lpcat3*^BKO^ mice acclimated to TN housing (30℃) for 2 weeks, and either kept at 30℃ or subjected to acute cold challenge (8℃) for 3 h (*n* = 4/group). (**D**) qPCR analysis of the thermogenic gene profiles in BAT from F/F and *Lpcat3*^BKO^ mice housed at 30℃ or subjected to acute cold challenge for 3 h (*n* = 4– 5/group). (**E**) Western blot analysis of phosphorylated PKA substrates in BAT from F/F and *Lpcat3*^BKO^ mice subjected to acute cold challenge (8℃) for 3 h (*n* = 4/group). GAPDH was used as a loading control. (**F**) Plasma NEFAs in F/F and *Lpcat3*^BKO^ mice housed at 30℃ or subjected to cold challenge (8℃) for 3 h (*n* = 4–13/group). (**G-I**) Seahorse respirometry performed on isolated BAT mitochondria from F/F and *Lpcat3*^BKO^ mice pre-acclimated to TN (*n* = 6–8/group). (**G**) CI/II–CIII–CIV activity measured in the presence of pyruvate (5 mM), malate (1 mM), ADP (4 mM), and GDP (1 mM). OCR traces (left) and quantification (right) of respiratory (3/4o/3u) states after serial injections with Oligo (4 μM), FCCP (4 μM), and Rot/AA (2/4 μM). (**H**) CI/II/ETF– CIII–CIV activity measured in the presence of malate (1 mM) and GDP (1 mM), using palmitoyl-carnitine as a substrate. OCR traces (left) and quantification (right) of respiratory (3/4o/3u) states after serial injections with PCarn/ADP, (40 μM/4 mM), Oligo (4 μM), FCCP (4 μM), and Rot/AA (2/4 μM). (**I**) CIV-specific activity measured in the presence of antimycin A (4 μM) and GDP (1 mM), using Asc/TMPD as substrates. OCR traces (left) and quantification (right) of CIV respiration after serial injections with Asc/TMPD (700 μM/500 μM) and sodium azide (33 mM). (**J**) Western blot analysis of OXPHOS subunits and UCP1 protein levels in isolated BAT mitochondria from F/F and *Lpcat3*^BKO^ mice kept at 30℃ (left) or subjected to acute cold challenge (8℃; right) for 3 h (*n* = 4/group). (**K**) BN–PAGE immunoblot analysis of isolated BAT mitochondria from F/F and *Lpcat3*^BKO^ mice subjected to acute cold challenge (8℃) for 3 h to assess SC assembly using OXPHOS complex-specific antibodies: CI (NDUFB8), CIII (UQCRFS1), or CIV (MTCO1 and COX4I1). SDS–PAGE Western blot analysis was performed on samples akin to those for BN-PAGE to serve as loading controls (*n* = 4/group). (**L**) BAT electron micrographs from F/F and *Lpcat3*^BKO^ mice subjected to acute cold challenge (8℃) for 3 h. (**M**) qPCR analysis of mtDNA copy number in BAT from F/F and *Lpcat3*^BKO^ mice housed at 30℃ or subjected to cold challenge (8℃) for 3 h (*n* = 5/group). (**N**) Electrophysiological recording of H^+^ currents across the IMM in BAT-derived mitoplasts from F/F and *Lpcat3*^BKO^ mice housed at 23℃. Top: voltage ramp protocol. Bottom: representative traces showing proton currents before (black) and after (red) addition of 1 mM GDP. Right: summary of H^+^ current amplitude (*n* = 9–11/group). All animals described in this Figure were 10-week-old males and pre-acclimated to TN (30℃) for 2 weeks, unless stated otherwise. Data are presented as mean ± SEM. \*\*\**p*<0.001, \*\*\*\**p*<0.0001 by two-sided Welch’s *t*-test (I, N) or one-way ANOVA with Tukey’s multiple comparisons test (D, F, G, H, M).

**Fig. S5.**
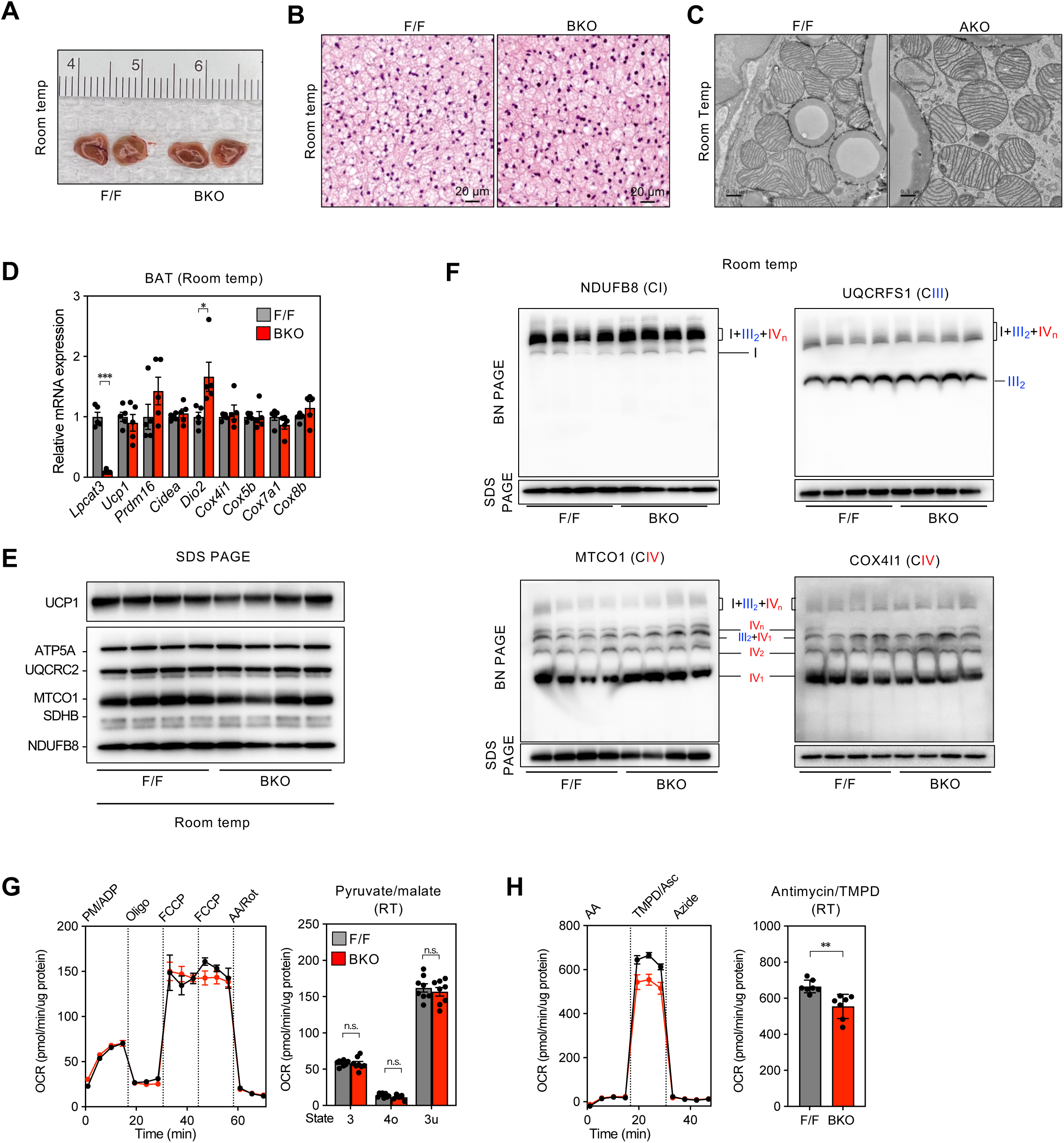
*Lpcat3*^BKO^ mice housed under standard laboratory conditions have no defects in BAT physiology. (**A-C**) Gross appearance (**A**), H&E stainings (**B**), and electron micrographs (**C**) of BAT from 10-week-old F/F and *Lpcat3*^BKO^ mice housed at room temperature (23℃). (**D**) qPCR analysis of thermogenic gene profiles in BAT from F/F and BKO mice housed at 23℃ (*n* = 5/group). (**E**) Western blot analysis of OXPHOS subunits and UCP1 protein levels in isolated BAT mitochondria from F/F and *Lpcat3*^BKO^ mice housed at 23℃ (*n* = 4/group). (**F**) BN–PAGE immunoblot analysis of isolated BAT mitochondria from F/F and *Lpcat3*^BKO^ mice housed at 23℃ to assess respiratory SC assembly using OXPHOS complex-specific antibodies: CI (NDUFB8), CIII (UQCRFS1), or CIV (MTCO1 and COX4I1). SDS–PAGE Western blot analysis was performed on samples akin to those for BN–PAGE to serve as loading controls (*n* = 4/group). SDS–PAGE images in (F) of NDUFB8 and MTCO1 are cropped images of (E). (**G, H**) Seahorse respirometry performed on isolated BAT mitochondria from F/F and *Lpcat3*^BKO^ mice housed at 23℃ (*n* = 7–8/group). (**G**) CI/II–CIII–CIV activity measured in the presence of pyruvate (5 mM), malate (1 mM), ADP (4 mM), and GDP (1 mM). OCR traces (left) and quantification (right) of respiratory (3/4o/3u) states after serial injections with Oligo (4 μM), FCCP (4 μM), and Rot/AA (2/4 μM). (**H**) CIV-specific activity measured in the presence of antimycin A (4 μM) and GDP (1 mM), using Asc/TMPD as substrates. OCR traces (left) and quantification (right) of CIV respiration following serial injections with Asc/TMPD (700 μM/500 μM) and sodium azide (33 mM). Data are presented as mean ± SEM. \**p*<0.05, \*\**p*<0.01, \*\*\**p*<0.001 by two-sided Welch’s *t*-test (D, G, H).

**Fig. S6.**
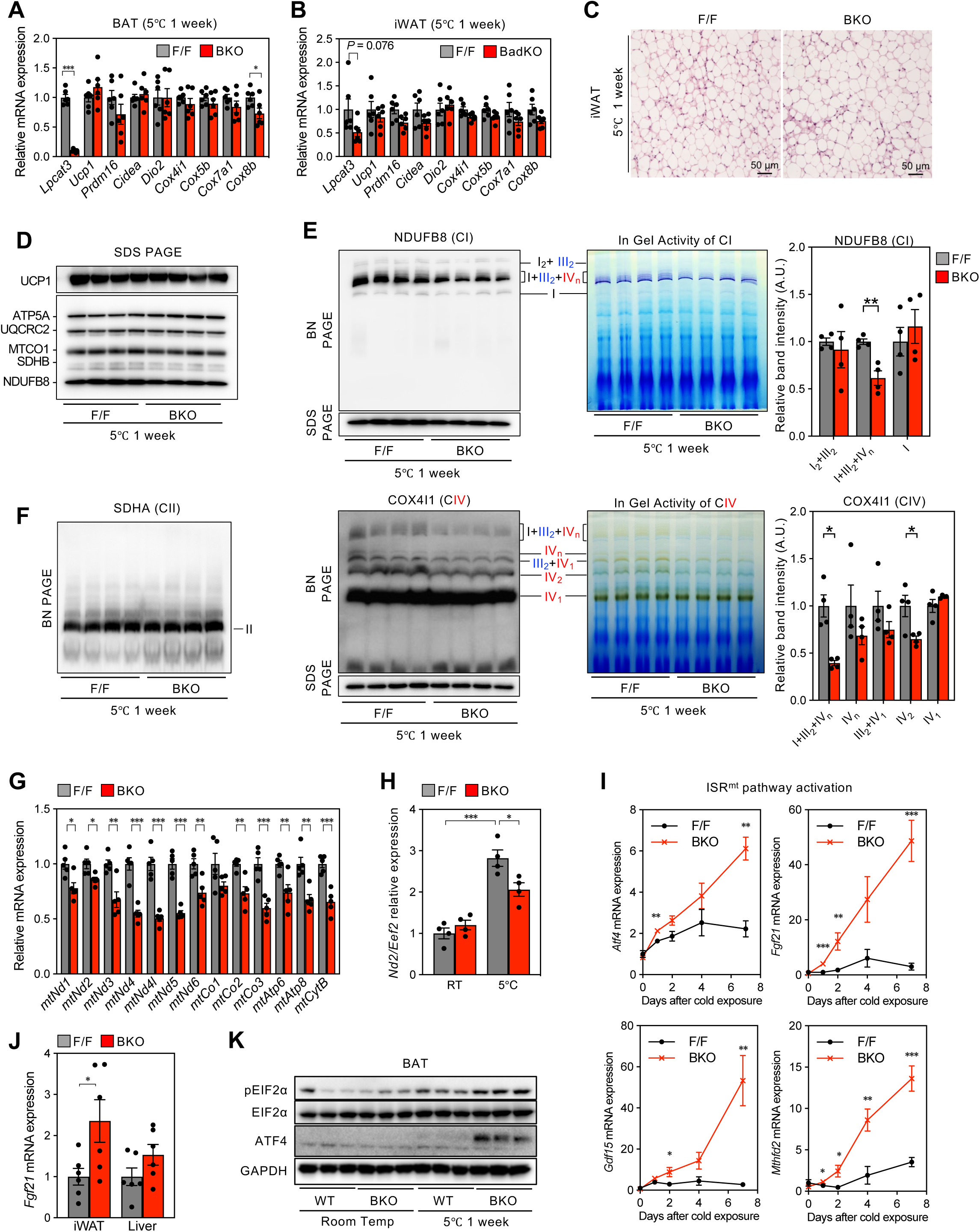
LPCAT3 deficiency in BAT leads to IRSmt activation during cold adaptation. (**A, B**) qPCR analysis of thermogenic gene profiles genes in (**A**) BAT and (**B**) iWAT from 10-week-old male F/F and *Lpcat3*^BKO^ mice acclimated to cold exposure (5℃) for 7 days (*n* = 6/group). (**C**) Representative H&E staining of iWAT sections from cold-acclimated F/F and *Lpcat3*^BKO^ mice. (**D**) Western blot analysis of OXPHOS subunits and UCP1 protein levels in cold-acclimated F/F and *Lpcat3*^BKO^ mice (*n* = 4/group). (**E**) BN–PAGE immunoblot analysis (left) of SC assembly in isolated BAT mitochondria from cold-acclimated F/F and *Lpcat3*^BKO^ mice, *in*–*gel* analysis of CI or CIV activity (middle) and quantification (right) using OXPHOS complex-specific antibodies for CI (NDUFB8) or CIV (COX4I1). SDS–PAGE Western blot analysis was performed on samples akin to those for BN–PAGE to serve as loading controls (*n* = 4/group). (**F**) BN–PAGE immunoblot analysis of CII (SDHA) in BAT mitochondria from cold-acclimated F/F and *Lpcat3*^BKO^ mice. (**G**) qPCR analysis of mtDNA-encoded gene transcripts in BAT from cold-acclimated F/F and *Lpcat3*^BKO^ mice (*n* = 5, 5). (**H**) qPCR analysis of mtDNA copy number in BAT from F/F and *Lpcat3*^BKO^ mice housed at room temperature (23℃) or acclimated to 5℃ for 7 days (*n* = 4/group). (**I**) qPCR analysis of ISR^mt^-related gene transcripts (*Atf4*, *Fgf21*, *Gdf15*, *Mthfd2*) in BAT from F/F and *Lpcat3*^BKO^ mice housed at (23℃) and subjected to cold exposure (5℃) for 0, 1, 2, 4, or 7 days (*n* = 4-5/group). (**J**) qPCR analysis of *Fgf21* gene transcript levels in iWAT and liver from F/F and *Lpcat3*^BKO^ mice housed at 23℃ or acclimated to 5℃ for 7 days (*n* = 6/group). (**K**) Western blot analysis of ISR^mt^ markers in BAT from F/F and *Lpcat3*^BKO^ mice housed at 23℃ or acclimated to 5℃ for 7 days (*n* = 3/group). All animals described in this Figure were 10-week-old males and reared at 23℃. Data are presented as mean ± SEM. \**p*<0.05, \*\**p*<0.01, \*\*\**p*<0.001 by two-sided Welch’s *t*-test (A, B, E, G, I, J), or one-way ANOVA with Tukey’s multiple comparisons test (H).

